# NaBC1 boron transporter enables myoblast response to substrate rigidity via fibronectin-binding integrins

**DOI:** 10.1101/2024.06.14.599051

**Authors:** Juan Gonzalez-Valdivieso, Giuseppe Ciccone, Eva Barcelona-Estaje, Aleixandre Rodrigo-Navarro, Rafael Castillo, Patricia Rico, Manuel Salmeron-Sanchez

## Abstract

Cells are sensitive to the physical properties of their microenvironment and transduce them into biochemical cues that trigger gene expression and alter cell behavior. Numerous proteins, including integrins, are involved in these mechanotransductive events. Here, we identify a novel role for the boron transporter NaBC1 as a mechanotransducer. We demonstrate that soluble boron ions activate NaBC1 to enhance cell adhesion and intracellular tension in C2C12 myoblasts seeded on fibronectin-functionalised polyacrylamide (PAAm) hydrogels. Retrograde actin flow and traction forces exerted by these cells are significantly increased *in vitro* in response to both increased boron concentration and hydrogel stiffness. These effects are fibronectin and NaBC1-mediated as they are abrogated in hydrogels coated with laminin-111 in place of fibronectin and in esiRNA NaBC1-silenced cells. Our findings thus demonstrate that NaBC1 controls boron homeostasis and also functions as a mechanosensor.

## Introduction

Cells sense mechanical signals from the surrounding environment, the extracellular matrix (ECM), and transduce them into biochemical signals through mechanotransductive processes ^1,2^. Whilst attaching to their underlying matrix, cells probe it to sense its stiffness, viscosity, and topography ^3–6^. Cells then bind to the ECM via integrins, which binds to ECM proteins through so-called focal adhesions (FA) ^7,8^. The molecular clutch model ^9–11^ explains how cells mechanically sense the ECM by an strengthening proteins involved in the mechanical links between the ECM, integrins and their actin cytoskeleton. These physical links involve other mechanosensitive proteins found in focal adhesions that can bind directly to actin, including talin and vinculin ^10,12–14^. The adhesion machinery these proteins create allows cells to exert forces on the substrate, which, in turn, allows them to migrate, proliferate, and even differentiate into multiple cell types ^4,15–19^. The molecular clutch is a well-accepted paradigm. However, recent studies suggest that other membrane proteins might play important roles in cell mechanotransduction ^20,21^ by helping to modulate the strength of cell adhesion and the transmission of forces between cells and the ECM, thereby enabling the dynamic regulation of cell behavior.

The NaBC1 boron (B) transporter is encoded by the *SLC4A11* gene and is a Na^+^-coupled B co-transporter that controls B homeostasis ^22^. Mutations in the *SLC4A11* gene are involved in rare diseases, such as endothelial corneal dystrophies ^23^. Previous studies have reported a role for B in osteogenic differentiation ^24^ and adipogenesis inhibition ^25^, however B function and homeostasis are not completely understood. We have previously demonstrated that NaBC1 crosstalk with growth factor receptors (GFR) enhances vascularisation ^26^, adhesion-driven osteogenesis ^27^, myogenic differentiation ^28^ and muscle regeneration *in vivo* ^29^.

In this work, we demonstrate that NaBC1 is a mechanosensitive protein, and that mechanotransduction happens via its interaction with fibronectin-binding integrins. We show that active-NaBC1 in C2C12 myoblasts enhanced cell spreading *in vitro* through the formation of more and larger focal adhesions in response to substrate stiffness. We also show that intracellular tension, cell stiffness, retrograde actin flow, and traction forces are upregulated by B in a substrate-stiffness-dependent manner. From our findings, we propose that NaBC1 is a mechanosensor that plays an important role in cell response to mechanical stimuli. This new role for NaBC1 might be important for understanding the pathologies that occur in skeletal muscle when cell-ECM membrane-cytoskeleton interactions are disrupted to cause muscular dystrophies ^30^.

## Results & Discussion

### Active-NaBC1 modulates cell response to substrate rigidity

PAAm hydrogels with tuneable properties are well-known systems that are widely used to study cell-ECM interactions to investigate processes such as cell migration, proliferation, malignancy, and differentiation ^31^. The stiffness of the ECM can activate different intracellular pathways and cytoskeletal arrangements, which modulate cell responses through the integrin receptors in the mechanotransductive process ^32^. Here, we investigate whether there are other, as yet unidentified, cell receptors and proteins that play a role in mechanotransduction.

In this study, we used PAAm hydrogels of different rigidities, as characterised by the Young’s modulus (*E*), which we termed soft (0.5 kPa), medium (9 kPa) and rigid (35 kPa) (Fig. 1A-B). First, PAAm hydrogels were functionalised with fibronectin to enable cell interactions, and then C2C12 myoblasts were seeded on top of them. We measured these cells’ viability and found it to be > 95% (Fig. S1A-B). The rigidity of the PAAm hydrogel substrate markedly influenced cell spreading and adhesion (Fig. 1C-D). On soft hydrogels, C2C12 myoblasts remained round in shape and did not properly attach to the substrate, but on medium and rigid substrates, they spread more and became larger. Substrate rigidity had an important effect on cell spreading since cell area significantly increased on the stiffest substrate (1515 ± 219 µm^2^), compared to cells seeded on medium hydrogels (1053 ± 154 µm^2^). We used concentrations of soluble B that have no effect on the viability of C2C12 myoblasts (Fig. S1C). Even though substrate rigidity had no effect on cell viability, cell proliferation was markedly altered (Fig. S1D). C2C12 myoblasts on soft hydrogels did not proliferate at a normal rate, relative to their proliferation on culture plates and on medium or rigid PAAm hydrogels. It is well known that, on soft substrates (0.5 kPa), adherent cells do not proliferate due to lack of adhesion ^33,34^.

**Figure 1.**
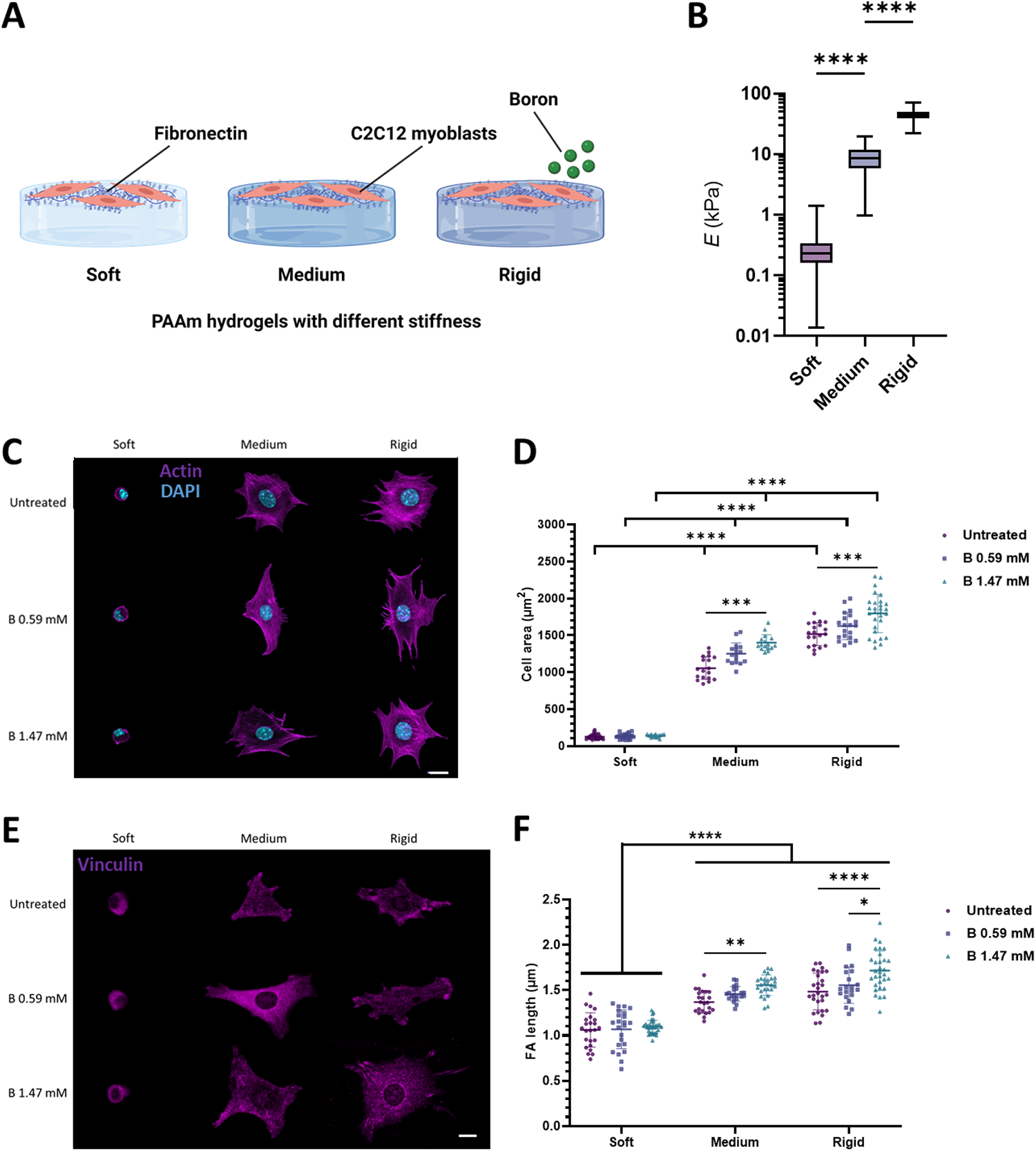
Active NaBC1 modulates cell response to substrate rigidity. A: Schematic representation of C2C12 myoblasts seeded on PAAm hydrogels with different mechanical properties (soft, medium and rigid) functionalised with fibronectin and treated with Boron (B). Scheme created with BioRender.com. B: Measurements of the elastic Young’s modulus of each hydrogel (soft, medium, rigid) by nanoindentation. *n* > 3 hydrogels with 9 repeated indentations on each single hydrogel. C: Representative immunofluorescence images of C2C12 myoblasts seeded on PAAm hydrogels of different stiffnesses, functionalised with fibronectin (FN) and stimulated with soluble B (0.59 and 1.47 mM). Magenta: actin cytoskeleton; Cyan: DAPI. Scale bar: 20 µm. D: Quantification of projected cell area of C2C12 myoblasts, cultured as described in panel B (0.59 and 1.47 mM). *n* = 10 cells from 3 different biological replicates. E: Representative immunofluorescence images of C2C12 myoblasts, cultured as described in panel B. Magenta: vinculin. Scale bar: 30 µm. F: Quantification of focal adhesion (FA) length in C2C12 myoblasts, cultured as described in panel B (0.59 and 1.47 mM). *n* = 10 cells from 3 different biological replicates. Data are represented as Mean ± Standard Deviation, and differences are considered significant for p ≤ 0.05 using one-way ANOVAs or two-way ANOVAs (Tukey’s multiple comparisons tests) for multiple comparisons. ***p ≤ 0.001, ****p ≤ 0.0001

On both medium and rigid substrates, we observed enhanced cell spreading (up to 1400 ± 110 and 1800 ± 260 µm^2^, respectively) when NaBC1 transporter was stimulated with soluble B, which was concentration-dependent in manner, as reported in previous studies conducted on glass ^29^.

Increased substrate rigidity induces the growth of FAs and increases intracellular tension, in agreement with the prediction of the molecular clutch model ^10,11,35^. We used vinculin to quantify the size of FAs. Both the size (Fig. 1E-F and S2B) and the number (Fig. S2A) of FAs significantly increased following NaBC1 stimulation (with 0.59 mM and 1.47 mM concentration of B) on medium and rigid substrates but not on soft ones. We note that these differences in cell adhesion to the substrate were due only to substrate rigidity and not to different densities of fibronectin on the PAAm hydrogels used (Fig. S3).

The Yes-associated protein (YAP) is a reporter for mechanotransduction ^14^. Phosphorylation of the myosin light chain (pMLC) regulates cytokinesis and plays an important role in diverse cell functions via a Rho-associated kinase (ROCK) pathway after the assembly of FAs ^36^. To determine whether the effect of NaBC1 on cell adhesion involves intracellular tension, we measured the phosphorylation of MLC (Fig. 2A-B) and YAP nuclear translocation (Fig. 2C-D). Our results show that both pMLC levels and YAP nuclear translocation were significantly increased on medium (up to 2.88 and 3.72-fold increase) and rigid (up to 2.79 and 5.98-fold increase) PAAm substrates after NaBC1 stimulation with soluble B. In cells seeded on soft gels, we observed no significant differences in pMLC levels nor in YAP nuclear translocation following NaBC1 stimulation, regardless of the concentration of B used.

**Figure 2.**
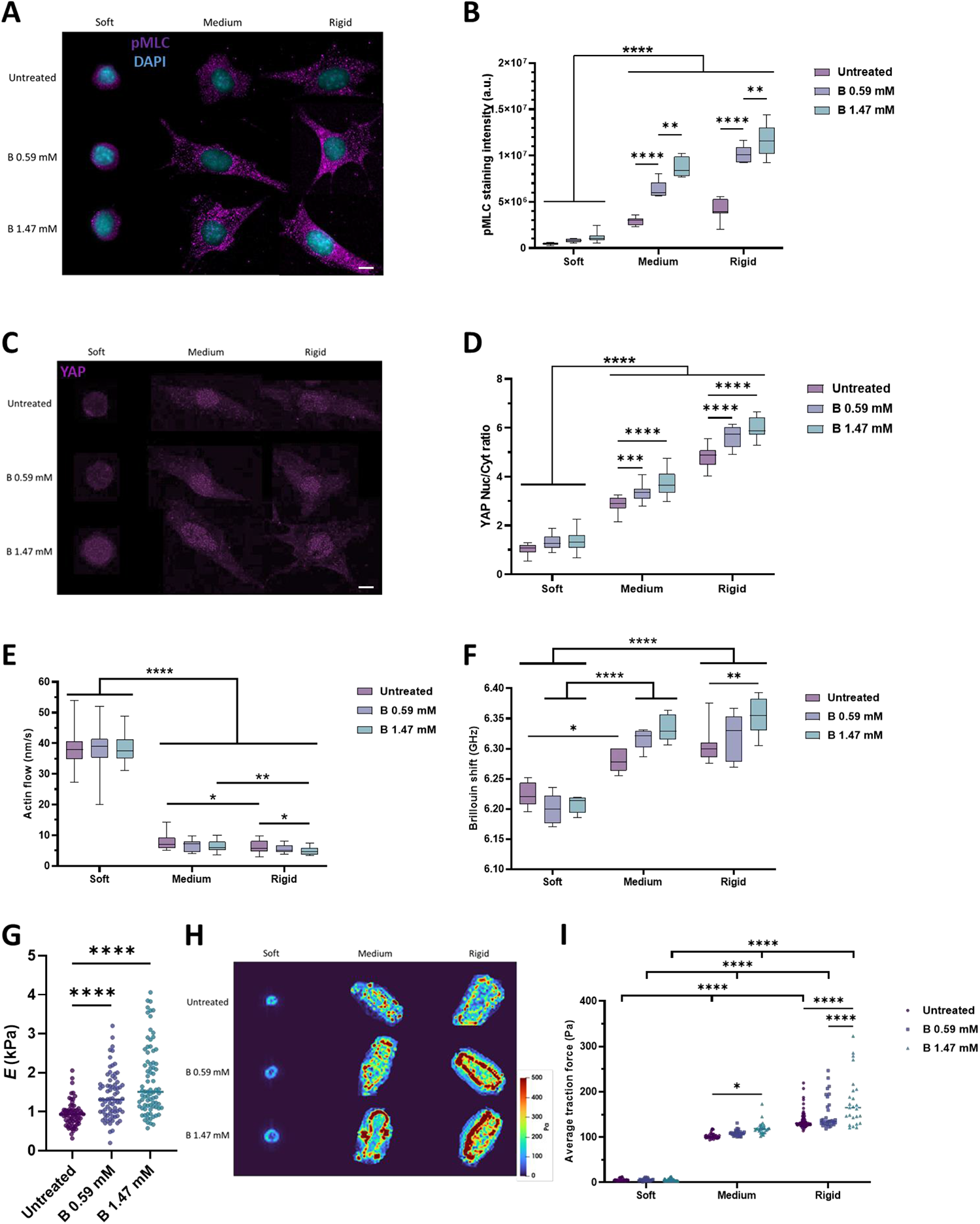
Active-NaBC1 enhances intracellular tension, retrograde actin flow and forces exerted by cells. The experimental results reported in A, B, C, D, E and F were obtained from culture conditions in which C2C12 myoblasts were seeded on PAAm hydrogels of different stiffnesses (soft, medium and rigid) that were functionalised with fibronectin (FN) and stimulated with soluble boron (B) (0.59 and 1.47 mM). A: Representative immunofluorescence images of C2C12 myoblasts cultured as described. Magenta: phosphorylated myosin light chain (pMLC); Cyan: DAPI. Scale bar: 50 µm. B: Quantification of pMLC intensity in C2C12 myoblasts cultured as described. *n* = 10 cells from 3 different biological replicates. C: Representative immunofluorescence images of C2C12 myoblasts cultured as described. Magenta: YAP. Scale bar: 50 µm. D: Quantification of YAP nuclear translocation in C2C12 myoblasts cultured as described. *n* = 10 cells from 3 different biological replicates. E: Quantification of actin retrograde flow in C2C12 myoblasts cultured as described. *n* = 5 cells with at least 5 different flow areas per cell. F: Quantification of Brillouin shift in C2C12 myoblasts cultured as described and imaged using Brillouin microscopy. *n* = 10 cells from 3 different biological replicates. G: Quantification of cell stiffness by nanoindentation of C2C12 myoblasts seeded on glass coverslips functionalized with FN and stimulated with soluble B (0.59 and 1.47 mM). *n* = 10 cells with 9 indentations on each single cell from 3 different biological replicates. H: Representative traction maps of C2C12 myoblasts cultured as described. I: Quantification of traction forces exerted by C2C12 myoblasts cultured as described. *n* = 30 cells from 10 different locations within each hydrogel from 3 different biological replicates. Data are represented as Mean ± Standard Deviation, and differences are considered significant for p ≤ 0.05 using one-way ANOVAs or two-way ANOVAs (Tukey’s multiple comparisons tests) for multiple comparisons. *p ≤ 0.05, **p ≤ 0.01, ***p ≤ 0.001, ****p ≤ 0.0001

The molecular clutch model links the retrograde flow of actin to the ECM through FAs and integrins. In this model, the elastic resistance of the substrate to deformation offsets the contractility of myosin, thereby slowing the actin flow and increasing the forces loaded onto integrins and FAs. The force loading rate increases with substrate rigidity ^9,10^. On soft substrates, the nucleus is relaxed, and cells present a poor cytoskeletal organisation and weakly assembled FAs (in which talin is folded and vinculin is not recruited), leading to a lack of force generation. As rigidity increases, the cells’ connection to the ECM becomes stronger, giving them time to load enough force onto the FAs. This force loading leads to the unfolding of talin and to the recruitment of vinculin, which enables the actin-FAs-ECM clutch. To test the role of NaBC1 in this context, actin flow was measured in live C2C12 myoblasts transfected with fluorescent actin. As expected, the retrograde actin flow decreased as substrate rigidity increased (Fig. 2E). Upon stimulation of NaBC1 with B, retrograde actin flow decreased in cells seeded on medium substrates (7.5 and 6.2 nm/s for 0.59 and 1.47 mM of B, respectively), and decreased further in myoblasts seeded on rigid substrates (5 and 4.8 nm/s for 0.59 and 1.47 mM, respectively) compared to cells on hydrogels of the same rigidity not treated with B. We propose that the slower actin flow observed following NaBC1’s stimulation with B involves an enhanced engagement of the molecular clutch, as supported by the enhanced cell adhesion recorded on medium and rigid PAAm hydrogels, and as shown in Figure 1. Together, these results suggest that NaBC1 might be linked to the molecular clutch.

We measured cell stiffness using Brillouin microscopy (Fig. 2F). This is a contact-free, label-free, non-invasive technique used to optically map the mechanical properties of biological materials ^37–40^. Brillouin microscopy has also been recently used to map the elastic properties of cancer cells on PAAm gels ^41^. Here, we used Brillouin microscopy to quantify the Brillouin shift (a proxy for elasticity) of C2C12 myoblasts on PAAm hydrogels – the larger the shift of the Brillouin peak, the higher the stiffness of the measured region ^42^. The resolution of this technique (< 1 µm), allows for the mapping of cell mechanical properties at several points. Cells on soft hydrogels showed a lower Brillouin shift (6.224 ± 0.021 GHz) compared to cells cultured on medium or rigid hydrogels (6.281 ± 0.019 and 6.305 ± 0.029 GHz, respectively). Interestingly, when NaBC1 was stimulated with 0.59 mM of B, the cells on medium (6.317 ± 0.018 GHz) and rigid hydrogels (6.321 ± 0.035 GHz) stiffened. This stiffening was more evident when C2C12 cells were stimulated with B at 1.47 mM, as the Brillouin shift increased to 6.334 ± 0.022 GHz and 6.353 ± 0.029 GHz on medium and rigid hydrogels, respectively (Fig. 2F and S4). We used nanoindentation to confirm the data obtained from Brillouin (Fig. 2G). C2C12 myoblasts on medium and soft hydrogels treated with B, either at 0.59 mM or 1.47 mM, presented with significantly higher Young’s modulus (1.31 ± 0.59 and 1.64 ± 0.89 kPa, respectively), compared to untreated cells (0.94 ± 0.31 kPa). Thus, higher cell stiffness is a consequence of NaBC1 stimulation as determined by two independent techniques and might be related to enhanced cell attachment.

Cells generate traction forces that deform the ECM and engage the molecular clutch. We therefore performed traction force microscopy (TFM) to measure cell forces on the different substrates and to measure changes in cell force after NaBC1 stimulation (Fig. 2H-I). We observed in our TFM results that traction stress gradually increased together with substrate rigidity. Stress maps indicate that higher forces are exerted on the cell edges, particularly on the stiffest surface (35 kPa), and consistent with the location of large FA complexes (Fig. 1 C-E). Our results also showed that, following NaBC1 stimulation, the lowest B concentration (0.59 mM) was sufficient to produce a significant increase in cell traction forces (151.1 ± 34.5 Pa) but only on the rigid substrate. When the concentration of B was increased to 1.47 mM, we observed increased cell traction forces on both the medium (119.8 ± 13.4 Pa) and rigid (178.9 ± 56.5 Pa) hydrogels. These results, together with the increased cell stiffness observed following B stimulation, indicate that higher intracellular tension occurs as a result of integrin and NaBC1 activation.

As explained above, cells seeded on soft PAAm hydrogels (0.5 kPa) adopted a different cell morphology to cells cultured on stiffer substrates, and showed different cell behaviors and proliferation rates as well. We therefore hypothesized that C2C12 myoblasts undergo cell senescence due to their lack of adhesion to their substrate. Cell senescence consists of a state in which cells remain metabolically active without undergoing cell death or division ^43^. It is involved in numerous biological processes, such as tumor suppression, tumor progression, aging, and tissue repair ^44^. Common markers of cell senescence include multinucleated cells, increased vacuolisation, morphological changes, and the expression of pH-dependent β-galactosidase ^45^. β-galactosidase resides in lysosomes and converts β-galactosides into monosaccharides under acidic pH. Its activity is 100% higher in senescent cells relative to pre-senescent cells ^46^. We therefore assayed β-gal activity to test our hypothesis that C2C12 myoblasts cultured on soft substrates undergo senescence (Fig. S5). Our results showed that the enzymatic activity of β-galactosidase was 5.82 and 5.08 times higher in C2C12 myoblasts seeded on soft hydrogels than in C2C12 myoblasts seeded on medium or rigid substrates, respectively. Interestingly, cell senescence decreased over time in C2C12 myoblasts seeded on soft hydrogels (Fig. S5A-B), which could be explained by ECM secretion from non-senescent cells. Moreover, NaBC1 stimulation with B had no effect on β-gal activity on any substrate, at early (Fig. S5C) nor long (Fig. S5D) time points. We can speculate that this arrest of the cell cycle might be responsible for the lack of response in cell mechanotransduction after NaBC1 stimulation.

### NaBC1 cooperates with fibronectin-binding integrins to modulate intracellular signalling

To decipher the role of NaBC1 in the molecular clutch, we investigated the PI3K/AKT signalling pathway as previous studies have highlighted that this pathway undergoes adhesion-dependent activation ^47^. Fig. 3A-D shows that *AKT* and *mTOR* gene expression levels were upregulated up to 4 fold in C2C12 myoblasts seeded on medium and rigid substrates (at 24 h and 96 h), compared to non-stimulated myoblasts, following NaBC1 stimulation. We have previously demonstrated that B-loaded hydrogels promote muscle regeneration *in vitro* and *in vivo* ^29^. We therefore assayed the expression of the Vascular Endothelial Growth Factor receptor (*VEGFR*), Insulin receptor (*INSR*), and insulin-like growth factor receptor (*IL-GFR*) genes, which are important for the functions of muscle cells ^48–50^. Our results showed that substrate rigidity had no influence on the expression of these genes. However, NaBC1 stimulation with B at 1.47 mM boosted *IL-GFR* gene expression up to 5 times on all substrates. Myogenin and myoD are typical markers of early myogenic differentiation ^51,52^. Fig. 3 shows that the expression of both markers was upregulated 4h after C2C12 myoblasts were seeded on medium substrates (9 kPa). These substrates have a similar level of elasticity to that of human healthy skeletal muscles, such as the *flexor digitorum profundus* (8.7 kPa) and the *gastrocnemius* (9.9 kPa) ^53^. By contrast to IL-GFR, which promotes muscle growth, growth differentiation factor 11 (GDF11) inhibits myogenesis via the phosphorylation of SMAD2/3 transcription factors ^54^. Our results show, for the first time, that NaBC1 stimulation downregulated *GDF11* expression in C2C12 myoblasts seeded on medium and rigid substrates after 24 hours, demonstrating a relationship between NaBC1 and GDF11 conditioned by the rigidity of the substrate.

**Figure 3.**
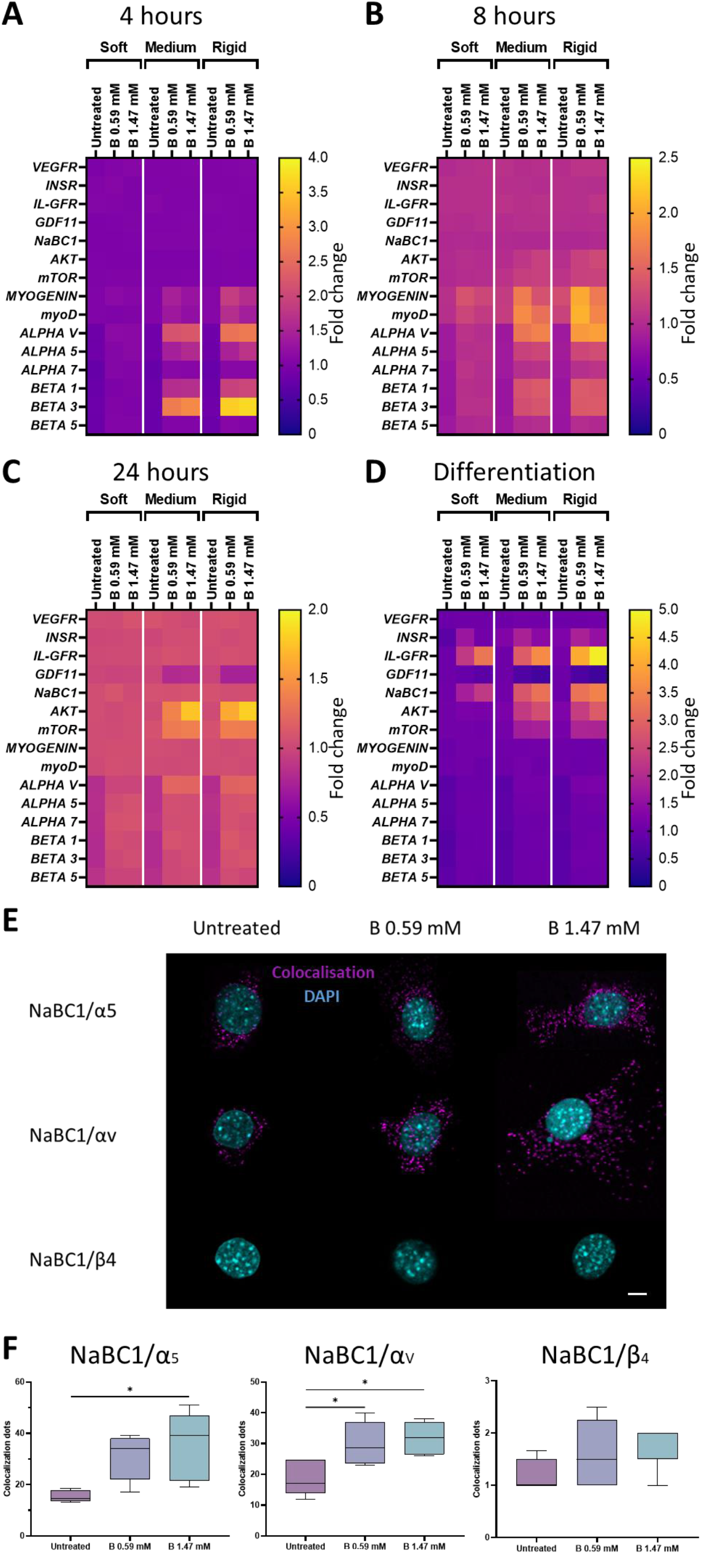
NaBC1 controls intracellular signalling via cooperation with fibronectin-binding integrins. A-C: Heat maps of gene expression of boron transporter (*NaBC1*), myogenesis markers (*MYOD, MYOGENIN*), AKT/mTOR pathway (*AKT, mTOR*), muscle metabolism (*IL-GFR*, *INSR*, *GDF11*, *VEGFR*), cell adhesion-related genes (*ALPHAV*, *ALPHA5*, *ALPHA7*, *BETA1*, *BETA3*, *BETA5* Integrins) in C2C12 myoblasts seeded on PAAm hydrogels of different stiffnesses, functionalized with fibronectin (FN) and stimulated with soluble boron (B) (at 0.59 and 1.47 mM) for 4 (A), 8 (B) or 24 hours (C) compared to untreated cells on cell culture plates, as measured by qPCR. D: Heat map of gene expression in C2C12 myoblasts seeded on PAAm hydrogels of different stiffnesses, functionalized with FN and stimulated with soluble B (at 0.59 and 1.47 mM) for 96 hours in myogenic differentiation conditions, as measured by qPCR. For figures A-D, *n* = 3 biological replicates with 3 technical replicates. E: Colocalization assays were performed by using the Duolink^®^ PLA protein detection technology, which is based on *in situ* proximity ligation assay (PLA) that allows the visualization and quantification of protein-protein interactions when proteins are present within 40 nm. Representative images showing the colocalization dots of NaBC1/α_5_, NaBC1/α_v_ and NaBC1/β_4_ in C2C12 myoblasts seeded on rigid PAAm hydrogels, functionalized with FN for 1 hour and stimulated with soluble B (0.59 and 1.47 mM). Magenta: colocalization dots; Cyan: DAPI. Scale bar: 50 µm. F: Quantification of number of colocalization dots of NaBC1/α_5_, NaBC1/α_v_ and NaBC1/β_4_. *n* = 30 cells from 3 different biological replicates. Data are represented as Mean ± Standard Deviation, and differences are considered significant for p ≤ 0.05 using one-way ANOVAs (Tukey’s multiple comparisons tests) for multiple comparisons. *p ≤ 0.05

Cells can perceive force through a variety of molecular sensors, of which ion-channels and transporters are the fastest and most efficient. Several studies have previously reported that ion-channels and integrins ^55^, as well as ion-transporters and integrins ^56^, can physically couple together to produce a cluster at the cell membrane. These clusters activate integrin-channel crosstalk and induce reciprocal signalling in which cell adhesion can induce channel activation ^57^ and channel engagement can regulate cell adhesion ^58,59^. Integrins might play a role in the localisation of ion-channels/transporters in the plasma membrane, and in channel regulation via the formation of macromolecular complexes that further regulate downstream signalling proteins. Indeed, ion channels sometimes transmit their signals through conformational coupling ^60^. The channel protein is not merely a final target, because it often feeds back by controlling integrin activation and/or expression, as occurs with different ion-channels that couple with integrin β_1_ and activate its expression ^61,62^. Here, we report that a combination of NaBC1 stimulation and substrate rigidity upregulates the expression of genes that encode fibronectin-binding integrins, such as α_v_, α_5_, β_1_ and β_3,_ in C2C12 myoblasts at 4 hours and, to a lesser extent, at 8 hours after NaBC1 stimulation (Fig. 4A-B). By contrast, the expression of α_7_ and β_5_ remained unaltered in C2C12 cells seeded on medium or rigid substrates. When C2C12 myoblasts were seeded on soft hydrogels, NaBC1 stimulation did not alter the downregulation of integrin expression (α_v_, α_5_, α_7_, β_1_, β_3_ and β_5_) (Fig. 4A-B), as is typical of substrates of low elastic moduli ^63^. To investigate whether integrin upregulation after NaBC1 stimulation occurred as a consequence of the interaction between α_5_β_1_ and α_v_β_3_ integrins and NaBC1, we measured the colocalisation of NaBC1/α_5_−α_v_ using the DUOLINK^®^ PLA kit system (Fig. 3E-F). Each dot in Fig 3E corresponds to the signal generated by two different proteins that are closer than 40 nm. Our results show that the addition of B to C2C12 myoblasts seeded on rigid substrates led to the colocalisation of fibronectin-binding integrins (α_5_ and α_v_) and NaCB1in a dose-dependent manner (Fig 3F). We did not observe any effect of B stimulation on the colocalisation of NaBC1 and other integrin receptors, such as integrin β_4_ (the laminin receptor) (Fig. 3E-F) ^64,65^. Together, these results demonstrate that NaBC1 cooperates with fibronectin-binding integrins in response to substrate rigidity. We therefore conclude that NaBC1 stimulation, in substrates of high enough elasticity, induces the expression of genes that encode fibronectin-binding integrins and components of the cell adhesion signalling pathways, AKT-mTOR.

**Figure 4.**
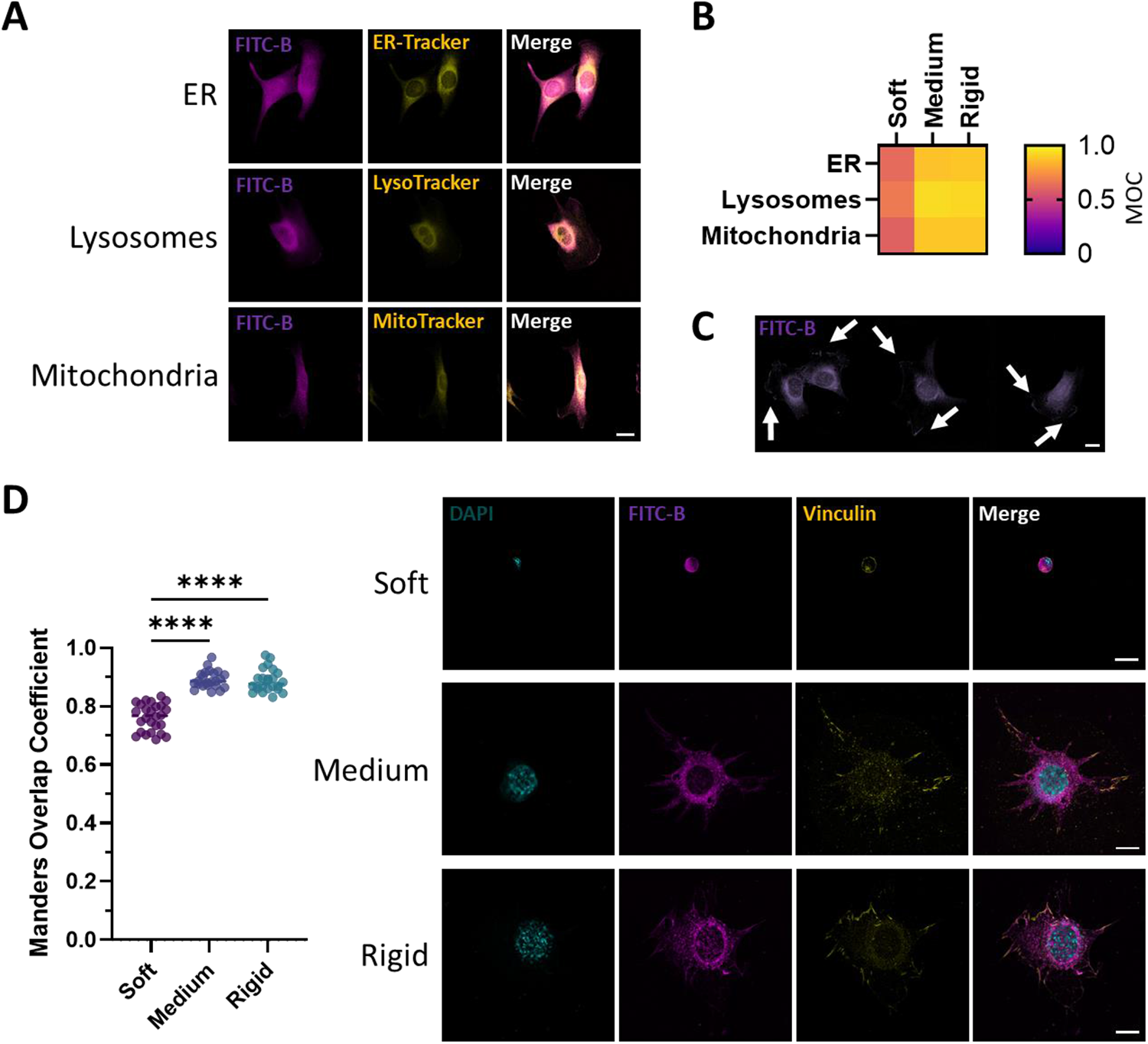
Boron subcellular localization highlights its presence in focal adhesions. A: Representative images of the colocalization of FITC-labelled Boron (FITC-B) and live trackers in C2C12 myoblasts seeded on rigid PAAm hydrogels functionalized with fibronectin (FN). Magenta: FITC-B. Yellow: live trackers for the: ER (ER-Tracker), lysosomes (LysoTracker) and mitochondria (MitoTracker). Scale bar: 20 µm. B: Heat map of B and live tracker colocalization in C2C12 myoblasts seeded on rigid PAAm hydrogels functionalized with FN. C: Representative images of the subcellular localization of FITC-B in C2C12 myoblasts seeded on rigid PAAm hydrogels functionalized with FN. Magenta: FITC-B. Arrows indicate clumps in cell edges. Scale bar: 20 µm. D: (Left) Quantification of FITC-B and focal adhesion colocalization in C2C12 cells seeded on hydrogels of different stiffnesses, as scored using the Manders Overlap Coefficient (MOC). (Right) Representative images of FITC-B and focal adhesion colocalization in C2C12 myoblasts seeded on rigid PAAm hydrogels functionalized with FN. Magenta: FITC-B; Yellow: vinculin; Cyan: DAPI. Scale bars: 20 µm. *n* = 20 cells from 3 different biological replicates. Data are represented as Mean ± Standard Deviation, and differences are considered significant for p ≤ 0.05 using one-way ANOVAs (Tukey’s multiple comparisons tests) for multiple comparisons. ****p ≤ 0.0001

### Intracellular dynamics of B

To test the intracellular dynamics of B, C2C12 myoblasts were incubated with FITC-labelled boron (FITC-B), in combination with multiple live trackers for specific cell compartments (the endoplasmic reticulum, lysosomes, and mitochondria). Fig. 4A-B shows that FITC-B colocalises with lysosomes, mitochondria, and the endoplasmic reticulum (ER) in C2C12 myoblasts seeded on medium and rigid hydrogels but not on soft substrates. We also observed FITC-B forming clumps at the cells’ edges (Fig. 4C). We therefore hypothesized that NaBC1 might be present in focal adhesions, since we had previously observed its colocalization with integrins. To test this hypothesis, we investigated the colocalization of FITC-B and vinculin (Fig. 4D). The Manders Overlap Coefficient (MOC) quantifies the amount of fluorescence overlaping between two channels ranging from 0 (an “anti-colocalization”) to 1 (a perfect colocalization) and is widely used as a quantitative tool to evaluate colocalization in biological microscopy ^66^. MOC was low for myoblasts seeded on soft hydrogels, but was close to 1 for those seeded on medium and rigid hydrogels. This result confirms the presence of NaBC1 in focal adhesions in cells seeded on medium/rigid substrates and corroborates the cooperation and colocalization between NaBC1 and fibronectin-binding integrins, triggered by the elasticity of the substrate.

FRAP (Fluorescence Recovery After Photobleaching) is a direct, non-invasive method used to study the mobility of biological molecules in living cells ^67^. FRAP corroborated our earlier finding that FITC-B colocalises with focal adhesions as demonstrated by the results we obtained with the living trackers (Fig. S6). Our FRAP results also showed that there were lower levels of FITC-B in the cell nuclei compared to the cytoplasm (Fig. S6A) and that FITC-B undergoes continuous influx into and efflux out of mitochondria and lysosomes (Fig. S6A). Furthermore, the short half-life and decreased fluorescence signal of FITC-B over time in focal adhesions indicate that FITC-B is internalized through these structures, supporting the colocalization of NaBC1 and fibronectin-binding integrins (Fig. S6B).

Given that FITC-B colocalises with mitochondria in C2C12 cells seeded on rigid substrates, we hypothesized that B might have a role in mitochondrial metabolism. Although previous studies have indicated that a relationship exists between cell mechanics and metabolism ^68^, how the mechanical cues exerted by the ECM influence metabolic pathways remains poorly understood ^69,70^. It is known that mechanotransduction pathways control cell processes (such as proliferation, differentiation, and death) that require energy generation and the biosynthesis of macromolecules ^71^. Indeed, approximately 50% of the ATP consumed by platelets ^72^ and neurons ^73^ is required to support the polymerization and rearrangement of the actin cytoskeleton. To test the effect of substrate elasticity and NaBC1 on cell metabolism, we measured total ATP and mitochondrial ATP (mATP) content in C2C12 myoblasts (Fig. S7). We detected the same level of total ATP for both medium and rigid substrates (Fig. S7A). When C2C12 myoblasts were seeded on rigid hydrogels, NaBC1 stimulation further increased their ATP content, relative to cells non treated with B on the same substrate, supporting a role for NaBC1 in mechanotransduction in the actin cytoskeleton of cells under higher tension (Fig. S7B) ^69,74^.

### NaBC1 is not involved in impaired cell response to stiffness on laminin-111-coated substrates

To test the role of NaBC1 in cell mechano-responses, we conducted further experiments on PAAm hydrogels coated with laminin-111 (Fig. 5) as laminin-111 has been recently shown to impair breast epithelial cell responses to substrate elasticity ^75^.

**Figure 5.**
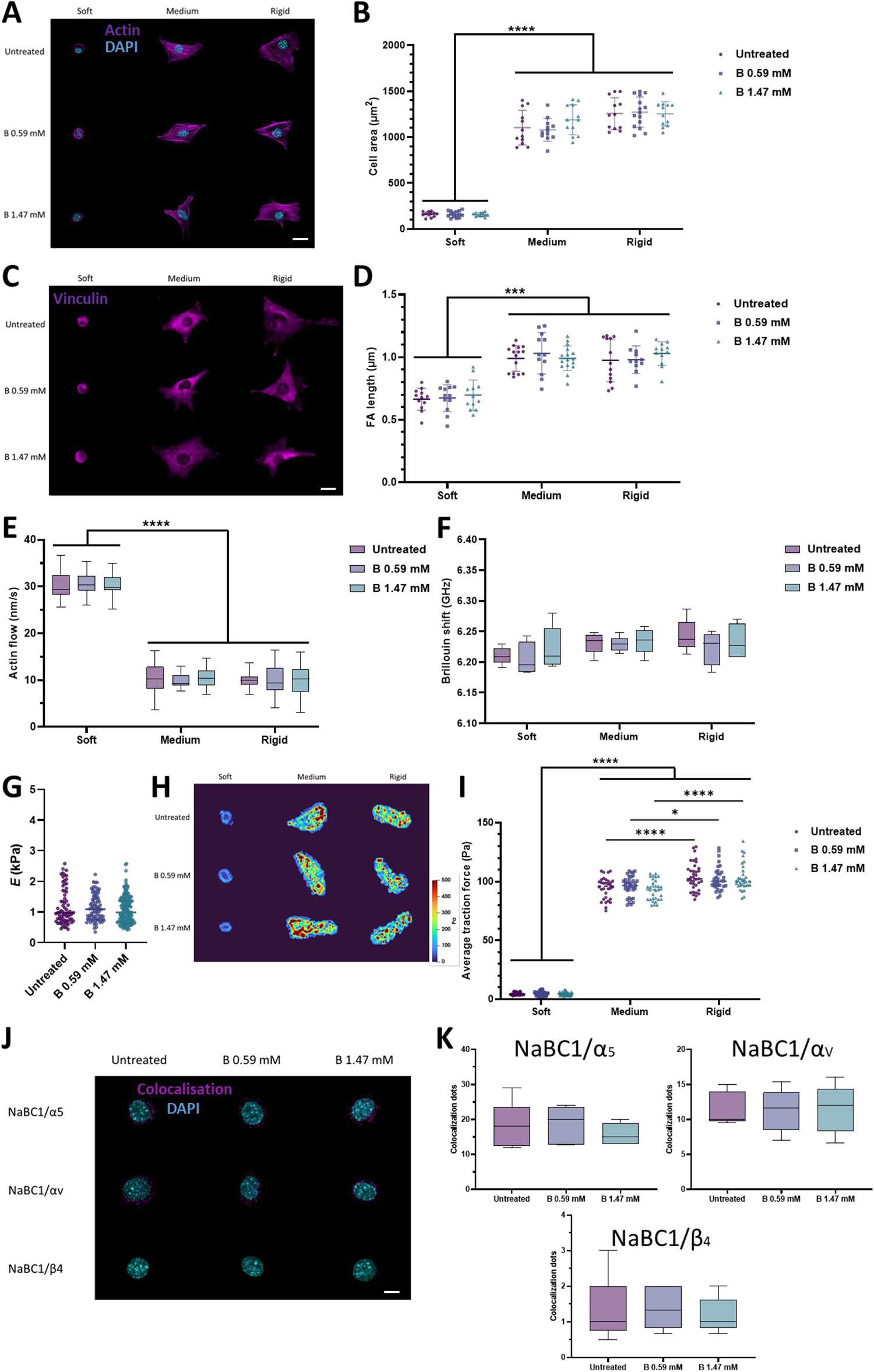
NaBC1 is not involved in impaired cell response to stiffness on laminin-111. The results reported in panels A-J derive from experiments in which C2C12 myoblasts were seeded on PAAm hydrogels of different stiffnesses (soft, medium, and rigid) that were functionalized with laminin-111 and stimulated with soluble boron ions at two different concentrations (0.59 and 1.47 mM). A: Representative immunofluorescence images of C2C12 myoblasts cultured as described. Magenta: actin cytoskeleton; Cyan: DAPI. Scale bar: 20 µm. B: Quantification of cell area of C2C12 myoblasts cultured as described. *n* = 10 cells from 3 different biological replicates. C: Representative immunofluorescence images of C2C12 myoblasts cultured as described. Magenta: vinculin. Scale bar: 20 µm. D: Quantification of focal adhesion (FA) length in C2C12 myoblasts cultured as described. *n* = 10 cells from 3 different biological replicates. E: Quantification of actin retrograde flow in C2C12 myoblasts cultured as described. *n* = 5 cells with at least 5 different flow areas per cell. F: Quantification of Brillouin shift in C2C12 myoblasts cultured as described and imaged with Brillouin microscopy. *n* = 10 cells from 3 different biological replicates. G: Quantification of cell stiffness by nanoindentation of C2C12 myoblasts seeded on glass coverslips functionalized with laminin-111 and stimulated with soluble B, as described. *n* = 10 cells with 9 indentations on each single cell from 3 different biological replicates. H: Representative traction maps of C2C12 myoblasts cultured as described. I: Quantification of traction forces exerted by C2C12 myoblasts when cultured as described. *n* = 30 cells from 10 different locations within each hydrogel from 3 different biological replicates. J: Colocalization assays were performed by using the Duolink^®^ PLA protein detection technology, which is based on *in situ* proximity ligation assay (PLA) that allows the visualization and quantification of protein-protein interactions when proteins are present within 40 nm. Representative images of colocalization dots of NaBC1/α_5_, NaBC1/α_v_ and NaBC1/β_4_ in C2C12 myoblasts cultured as described. Magenta: colocalization dots; Cyan: DAPI. Scale bar: 50 µm. K: Quantification of number of colocalization dots of NaBC1/α_5_, NaBC1/α_v_ and NaBC1/β_4_. *n* = 30 cells from 3 different biological replicates. Data are represented as Mean ± Standard Deviation, and differences are considered significant for p ≤ 0.05 using one-way ANOVAs or two-way ANOVAs (Tukey’s multiple comparisons tests) for multiple comparisons. *p ≤ 0.05, ***p ≤ 0.001, ****p ≤ 0.0001

We observed that when C2C12 cells were seeded onto soft PAAm hydrogels coated with laminin-111, they remained small and did not spread (Fig 5). We observed the same response in C2C12 cells seeded on medium and rigid hydrogels coated with laminin-111. This is in contrast to what we had previously observed: i.e., a continuous increase in cell area when cells were seeded on fibronectin-coated hydrogels, as the rigidity of the substrate increased (Figure 1). Importantly, in substrates coated with laminin-111, NaBC1 stimulation with soluble B at different concentrations did not alter this rounded cell morphology, as reported earlier in Fig 1, and the cell adhesive response on rigid and medium substrates was similar between them.

When C2C12 myoblasts were seeded on soft hydrogels coated with laminin-111, they were lower in number and formed small FAs (Fig. 5D, S8, S19, S20 and Fig. S21). These cells were also unable to respond to substrate stiffness, and differences in FA length and cell number between medium and rigid hydrogels were not observed (∼30 FA and 0.4 µm). Interestingly, B did not increase the length of FAs on these substrates, in contrast to what we had previously observed in cells seeded on fibronectin-coated substrates (Fig 1). We also assessed retrograde actin flow (Fig 5E) and observed the same trend: that the previously observed difference in retrograde actin flow in cells cultured on medium and stiff fibronectin-coated substrates was lost when cells were seeded on medium and stiff laminin-111 coated substrates. We also saw no retrograde actin flow response to the stimulation of NaCB1 with B in these cells (Fig 5E). Cell stiffness and cell traction forces exerted on laminin-111-functionalized substrates were measured using Brillouin microscopy/nanoindentation and traction force microscopy (Fig 5F-I). Undoubtedly, the weak adhesion of cells to laminin-111-coated substrates might explain the higher retrograde actin flow, lower cell stiffness and lower forces exerted, in comparison to cells on fibronectin-coated substrates. We also performed PLA colocalization assays and observed no interactions between NaBC1 and fibronectin-binding (α_5_−α_v_) or laminin-binding (β_4_) integrins on PAAm hydrogels of different stiffnesses that were functionalized with laminin-111 (Fig. 5J-K). The ATP content (both total and mATP) remained constant in cells on medium and rigid substrates even after incubation with soluble B (Fig. S10). The lack of response to the B-mediated stimulation of NaCB1 on laminin-coated substrates – which hinder mechanotransduction and do not engage the molecular clutch ^75^ – support the mechanosensitive nature of NaCB1.

### NaBC1-silencing decreases cell mechanotransduction on fibronectin-coated surfaces

To demonstrate that the enhanced mechanotransductive effect of B on fibronectin-coated substrates happens through NaBC1, we silenced NaCB1 using a NaCB1-targeting esiRNA in C2C12 myoblasts (Fig. S11). esiRNA are comprised of a heterogeneous pool of siRNA all targeting the same mRNA sequence, thereby allowing us to perform post-transcriptional silencing of NaBC1 in a highly specific and effective gene knockdown without off-target effects. Fig. S11B shows that esiRNA-mediated silencing reduced NaBC1 expression to less than 10% of that in wild-type myoblasts but did not affect cell viability (Fig. S11C). Furthermore, Fig. S12 shows that cell transfection with esiRNA Control did not alter the cell response to substrate stiffness nor NaBC1 stimulation with soluble B. However, cell area and FAs were smaller and less numerous in NaCB1-silenced cells, relative to wild-type and transfection controls, indicative of lower cell spreading and poor adhesion (Fig. 6A-B, S14, S19, S20 and S21). Following NaCB1 silencing, C2C12 cells were incubated with soluble B. The cell area and FA length of these cells remained the same as that of untreated cells, indicating that the NaBC1 receptor mediates the effects of B on the adhesiveness and morphology of these cells. We next measured retrograde actin flow in NaBC1-silenced myoblasts, and observed no significant differences in retrograde actin flow between cells cultured on a medium or rigid substrate (Fig. 6C). When NaBC1 was stimulated with soluble B in NaBC1-silenced cells, it did not trigger any further decrease in actin flow. Unexpectedly, in NaBC1-silenced cells after incubation with soluble B, we observed an increased Brillouin shift (up to 6.322 ± 0.018 GHz) (as measured by Brillouin microscopy, Fig. 6D and S15) and Younǵs modulus measured by nanoindentation (up to 1.21 ± 0.48 kPa) (Fig. 6E). These increases were also noticeable in C2C12 myoblasts transfected with esiRNA Control (Fig. S12). However, untreated NaBC1-silenced myoblasts presented similar Younǵs modulus to Control-silenced myoblasts and were 24% softer than untreated wild-type myoblasts (Fig. 6F) suggesting that the silencing process has an effect in cell membrane and stiffness. Since the silencing process inherently cannot achieve 100% efficiency, the increase in cell stiffness observed in NaBC1-silenced cells following B stimulation may be attributed to the residual low levels of NaBC1 expression (<10% of the wild-type levels) (Fig. S11). TFM measurements show that NaBC1-silenced myoblasts exerted 20% lower forces, compared to wild-type cells, and that this was not increased by B stimulation (Fig. 6G-H). These decreased traction forces were not showed by myoblasts transfected with esiRNA Control (Fig. S12), thereby discarding any effect of cell transfection. The total and mATP content also remained unaltered following the addition of B to NaBC1-silenced cells (Fig. S16). Together, our results demonstrate that NaBC1 is responsible for enhanced mechanotransduction on fibronectin-coated substrates after B stimulation of cells.

**Figure 6.**
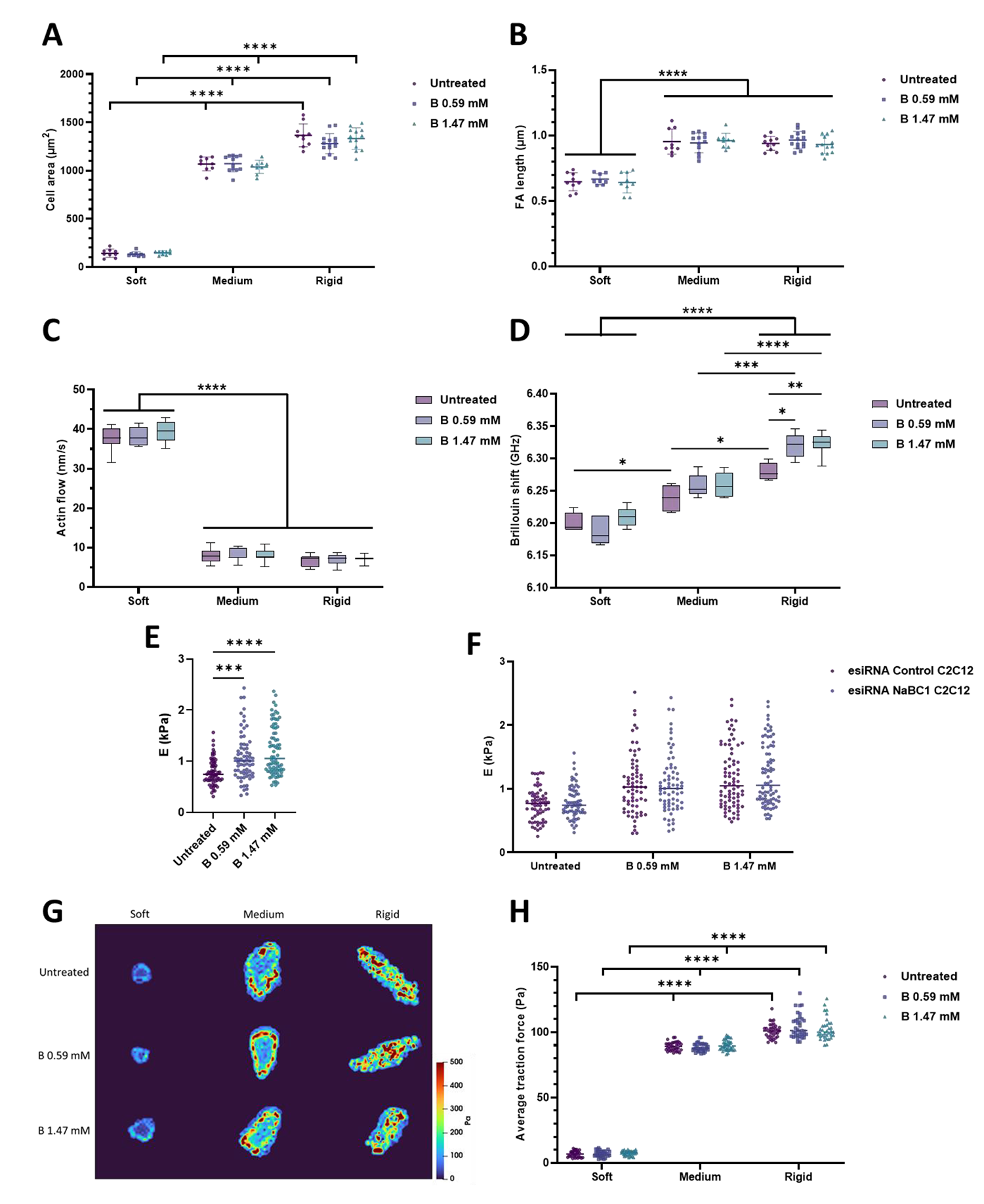
Cell mechanotransduction on fibronectin-coated surfaces is dependent of NaBC1. The results reported in panels A-D derive from experiments in which NaBC1-silenced C2C12 myoblasts were seeded on PAAm hydrogels of different stiffnesses (soft, medium, and rigid) that were functionalized with fibronectin (FN) and stimulated with soluble boron ions (B) at two different concentrations (0.59 and 1.47 mM). A: Quantification of cell area of NaBC1-silenced C2C12 myoblasts that were treated and cultured as described. *n* = 10 cells from 3 different biological replicates. B: Quantification of focal adhesion (FA) length in NaBC1-silenced C2C12 myoblasts that were treated and cultured as described. *n* = 10 cells from 3 different biological replicates. C: Quantification of actin retrograde flow in NaBC1-silenced C2C12 myoblasts that were treated and cultured as described. *n* = 5 cells with at least 5 different flow areas per cell. D: Quantification of Brillouin shift in NaBC1-silenced C2C12 myoblasts that were treated and cultured as described and imaged by Brillouin microscopy. *n* = 10 cells from 3 different biological replicates. E: Quantification of cell stiffness by nanoindentation of NaBC1-silenced C2C12 myoblasts seeded on glass coverslips functionalized with FN and stimulated with soluble B (0.59 and 1.47 mM). *n* = 10 cells with 9 indentations on each single cell from 3 different biological replicates. F: Comparison of cell stiffness by nanoindentation of NaBC1-silenced and Control-silenced C2C12 myoblasts seeded on glass coverslips functionalized with FN and stimulated with soluble B (0.59 and 1.47 mM). *n* = 10 cells with 9 indentations on each single cell from 3 different biological replicates. G: Representative traction maps of NaBC1-silenced C2C12 myoblasts that were treated and cultured as described. H: Quantification of traction forces exerted by NaBC1-silenced C2C12 myoblasts that were treated and cultured as described. *n* = 30 cells from 10 different locations within each hydrogel from 3 different biological replicates. Data are represented as Mean ± Standard Deviation, and differences are considered significant for p ≤ 0.05 using one-way ANOVAs or two-way ANOVAs (Tukey’s multiple comparisons tests) for multiple comparisons. *P ≤ 0.05, **P ≤ 0.01, ***P ≤ 0.001, ****P ≤ 0.0001

### The influence of NaBC1 on the dynamics of the actin cytoskeleton is linked to talin-vinculin binding

We performed experiments to investigate potential crosstalk between the NaBC1 receptor and the molecular clutch. To do so, we transfected C2C12 myoblasts with the VD1 plasmid (Fig. 7 and S17), which encodes a dominant protein composed of the head domain of vinculin that out competes endogenous vinculin for talin binding ^6,10,76^. Thus, VD1-transfected cells break the link between integrins and the actin cytoskeleton, preventing the cells’ response to stiffness, which is mediated by talin’s unfolding ^77^. Fig. 7B shows that the cell area of VD1-transfected cells remains different on hydrogels of increasing stiffness (1047.16 ± 70.62 and 1364.78 ± 188.79 µm^2^ on medium and rigid hydrogels, respectively). When NaBC1 was activated in VD1 cells with different concentrations of B, cell area increased (Fig. 7B), reaching values that are higher than those observed in wild-type myoblasts (Fig. 1D, 1274.40 ± 162.50 and 1539.66 ± 151.31 µm^2^ on medium and rigid hydrogels after stimulation with B 1.47 mM, respectively).

**Figure 7.**
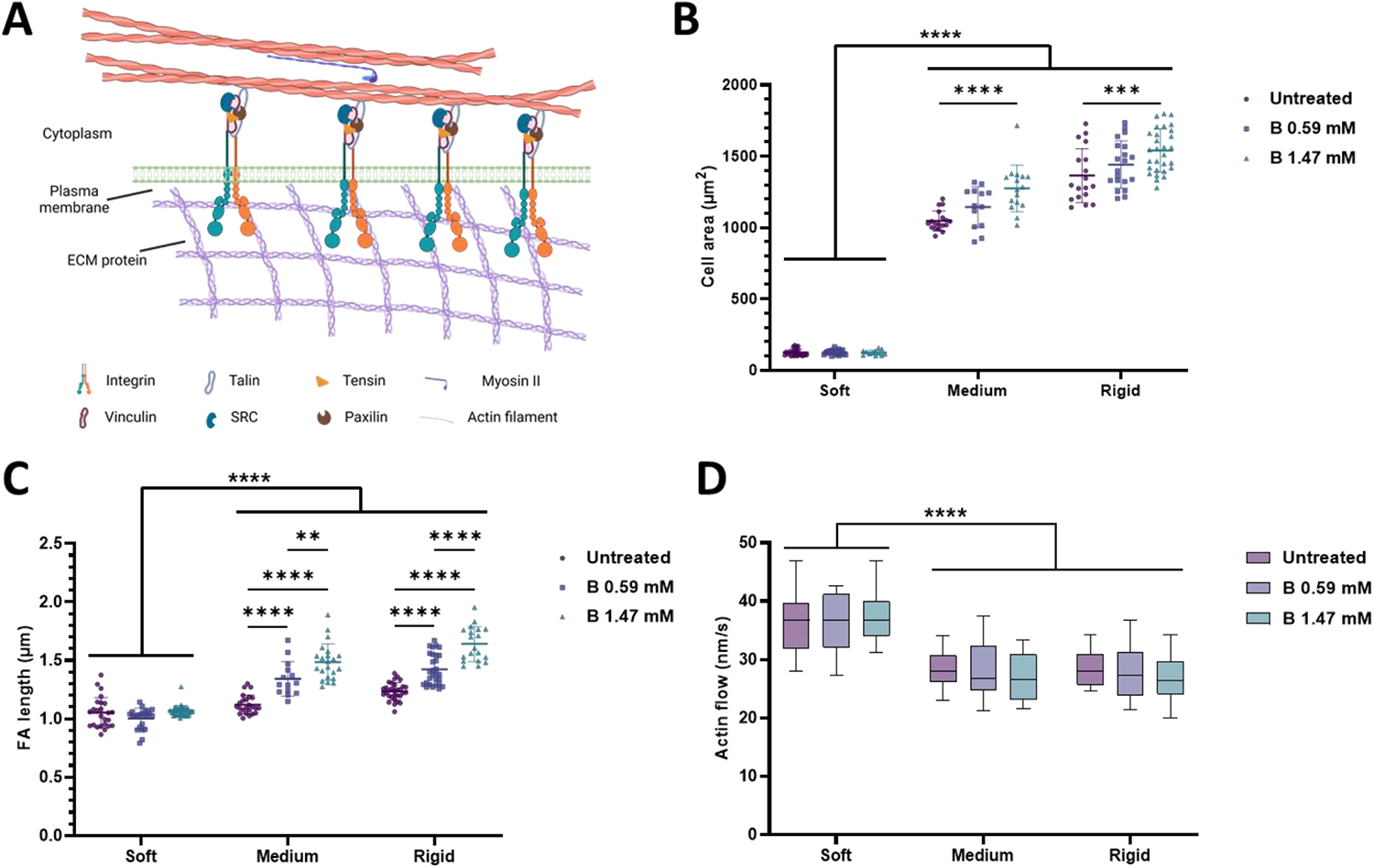
NaBC1 functions as a mechanosensor and is dependent of talin-vinculin binding. A: Schematic representation of the adhesion of a cell to ECM proteins via integrins. When talin unfolds in response to substrate rigidity vinculin can be recruited together with other proteins involved in the molecular clutch, forming mature focal adhesions and establishing the actin cytoskeleton. Schematic created with BioRender.com. Transfection with the VD1 plasmid, which encodes the vinculin head domain that can dominantly bind talin over endogenous vinculin, prevents the link between integrins and the actin cytoskeleton from being formed. Thus, VD1 mutant prevents cells’ response to the dynamics of the actin cytoskeleton that is mediated by talin unfolding. B: Quantification of cell area of C2C12 myoblasts transfected with VD1 seeded on PAAm hydrogels of different stiffnesses, functionalized with fibronectin (FN) and stimulated with soluble boron (B) (0.59 and 1.47 mM). *n* = 10 cells from 3 different biological replicates. C: Quantification of focal adhesion (FA) length in C2C12 myoblasts treated and cultured as described in panel B. *n* = 10 cells from 3 different biological replicates. D: Quantification of actin retrograde flow in C2C12 myoblasts treated and cultured as described in panel B. *n* = 5 cells with at least 5 different flow areas per cell. Data are represented as Mean ± Standard Deviation, and differences are considered significant for p ≤ 0.05 using two-way ANOVAs (Tukey’s multiple comparisons tests) for multiple comparisons. ***p ≤ 0.001, ****p ≤ 0.0001

The transfection of C2C12 myoblasts with VD1 did not alter the formation of FAs, neither their length (Fig. 7C) nor number (Fig. S18) compared to wild-type cells. However, on stimulation of these cells with B, the number and size of their FAs increased (Fig. 7C, S19, S20 and S21). VD1 impairs the link between integrins and the actin cytoskeleton and, as expected, transfected myoblasts with VD1 presented with high actin flow on medium (28.2 ± 2.8 nm/s) and rigid (28.4 ± 2.8 nm/s) hydrogels (Fig. 7D), close to the actin flow values recorded for cells on soft substrates and significantly different from those of wild-type myoblasts on hydrogels coated with fibronectin. The incubation of VD1 transfected cells with soluble B did not change their actin flow rates significantly, suggesting that the effect of NaBC1 on the dynamics of the actin cytoskeleton are dependent of proper talin folding/unfolding. We therefore conclude that our results and the molecular clutch model explain the active role of the NaBC1 transporter as a mechanosensor in response to stiffness.

### NaBC1 stimulation induces myogenic differentiation in medium and rigid substrates

To understand the biological importance of the NaBC1 B transporter, we studied the formation of myotubes *in vitro* by determining the expression of sarcomeric α-actinin, a typical marker for myogenesis ^78^. The fusion of myoblasts into myotubes is a phase of skeletal myogenesis that is essential for muscle repair ^79,80^. After cell adhesion, when cultured in low serum conditions, myoblasts spread, elongate, and fuse into myotubes ^81,82^. Figure 8 shows C2C12 myotubes after 4 days of culture under differentiation conditions (no serum and 1% insulin-transferrin-selenium (ITS)). When C2C12 myoblasts were cultured on fibronectin-coated hydrogels, NaBC1 stimulation with soluble B significantly induced myogenic differentiation on both medium and rigid substrates in a dose-dependent manner (25.5 ± 2.9 % and 33.8 ± 3.9 % on medium hydrogels vs. 21.5 ± 3.6 % and 27.8 ± 4.2 % on rigid hydrogels after incubation with 0.59 and 1.47 mM, respectively). Interestingly, the highest level of myogenic differentiation was achieved on medium hydrogels (9 kPa), which have a stiffness similar to that of healthy human skeletal muscles, such as the *flexor digitorum profundus* (8.7 kPa) and the *gastrocnemius* (9.9 kPa) ^53^. The enhanced myogenic differentiation mediated by B stimulation of NaBC1 was not observed in C2C12 myotubes cultured on laminin-111-coated hydrogels, which demonstrates the importance of mechanotranduction in NaCB1-treated cells which does not occur on cells cultured on laminin-111 coated hydrogels (Fig 5). Importantly, the stimulation of esiRNA-silenced NaBC1 cells with B also did not increase the percentage of differentiated cells. Finally, in C2C12 myoblasts transfected with the VD1 plasmid, we observed a dose-dependent percentage of differentiation after NaBC1 stimulation (up to 15.3 %) but markedly lower compared to wild-type myoblasts (up to 33.8 %). Together, our results show that the stimulation of NaBC1 with B in C2C12 myotubes cultured on medium or rigid substrates induces myogenic differentiation through a mechanism that involves cooperation with the molecular clutch through fibronectin-binding integrins.

**Figure 8.**
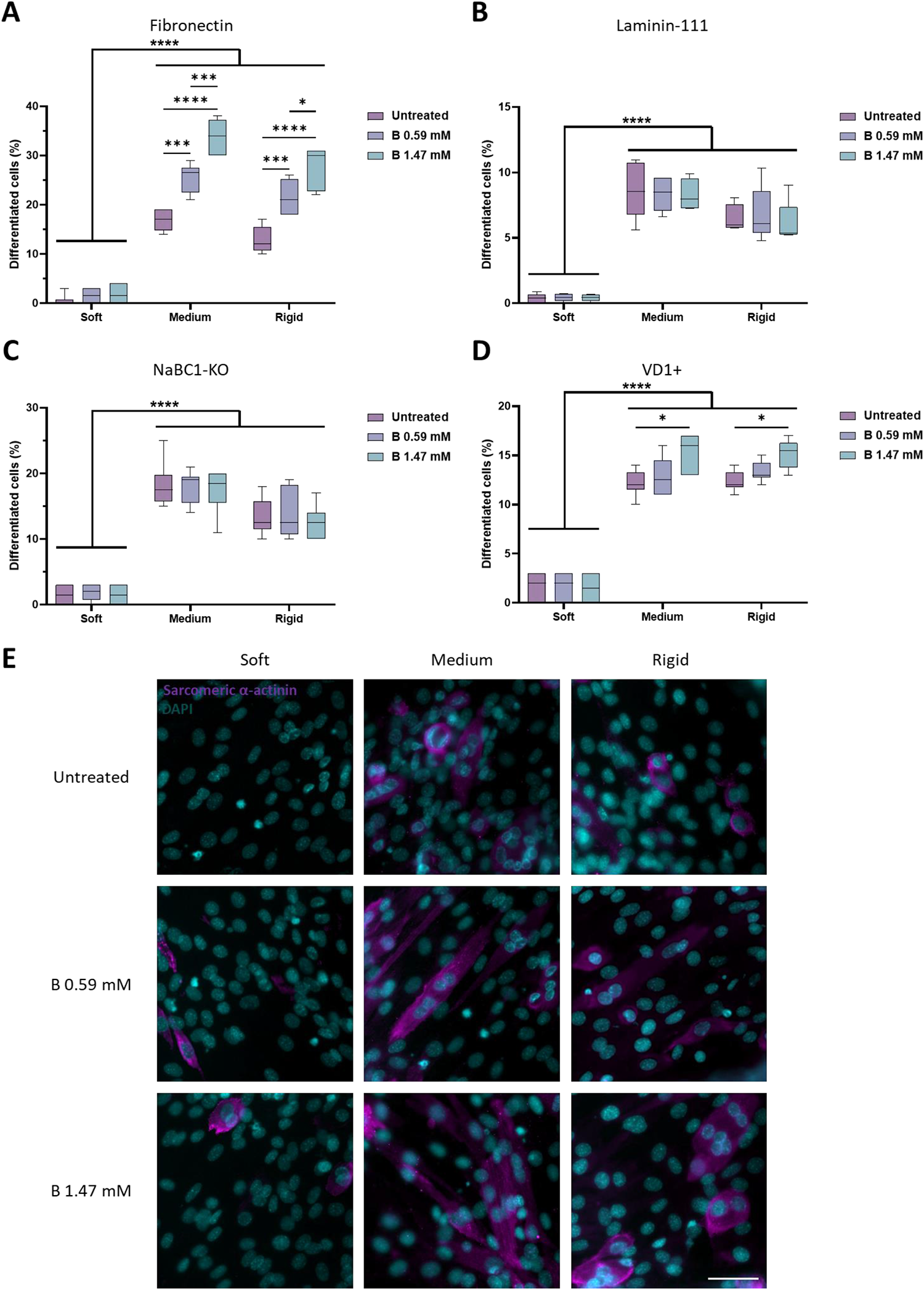
NaBC1 induces myogenic differentiation on fibronectin-coated medium and stiff substrates. A: Quantification of myogenic differentiation in C2C12 myoblasts seeded on PAAm hydrogels of different stiffnesses, functionalized with fibronectin (FN) and stimulated with soluble boron (B) (0.59 and 1.47 mM). *n* = 10 images from 3 different biological replicates. B: Quantification of myogenic differentiation in C2C12 myoblasts seeded on PAAm hydrogels of different stiffnesses, functionalized with laminin-111 and stimulated with soluble B ions (0.59 and 1.47 mM). *n* = 10 images from 3 different biological replicates. C: Quantification of myogenic differentiation in NaBC1-KO C2C12 myoblasts, cultured as described in A. *n* = 10 images from 3 different biological replicates. D: Quantification of myogenic differentiation in C2C12 myoblasts transfected with the VD1 plasmid and cultured as described in A. *n* = 10 images from 3 different biological replicates. E: Representative images of myogenic differentiation in C2C12 myoblasts, cultured as described in A. Magenta: sarcomeric α-actinin; Cyan: DAPI. Scale bar: 100 µm. Myotubes were counted when three or more cell nuclei were aligned. Data are represented as Mean ± Standard Deviation, and differences are considered significant for p ≤ 0.05 using two-way ANOVAs (Tukey’s multiple comparisons tests) for multiple comparisons. *p ≤ 0.05, ***p ≤ 0.001, ****p ≤ 0.0001

## Conclusion

We show that the NaBC1 transporter is actively involved in regulating cell behavior and fate beyond its role in B homeostasis. We demonstrated that the response of myoblasts to substrates stiffnesses which has been described by the molecular clutch model is altered by stimulation of the NaBC1 boron transporter. NaBC1 plays an important role in cell mechanotransduction and contributes to cell response to substrate rigidity in coordination with fibronectin-binding integrins (α_v_, α_5_, β_1_, and β_3_). Importantly, we show the absence of a response to substrate stiffness in NaBC1-silenced cells. Further, that NaBC1 stimulation had no effect on cell adhesion in myoblasts cultured on laminin-111-coated substrates, on which the cellular response to substrate rigidity is impaired and independent of the actin-talin-integrin molecular clutch ^75^.

### Experimental procedures Preparation of PAAm hydrogels

For PAAm hydrogels, all reagents were acquired from Sigma-Aldrich. Briefly, 1 mL volumes were prepared using stock solutions of 40% acrylamide (AAm) and 2% N,N’-methylenebisacrylamide (BisAAm) mixed in different ratios for specific hydrogel stiffnesses (Table 1). Solution volumes were then made up to 400 µL with milli-Q water, 25 µL 1.5% (w/w) tetramethylethylenediamine (TEMED) and 8 µL 5% (w/w) ammonium persulfate (APS) and mixed thoroughly. 10 µL of solution was spotted onto hydrophobic glass slides before placing acrylsilanized glass coverslips onto the spots. Gelation was allowed to occur at room temperature for 30 min before detaching and swelling in milli-Q water overnight at 4°C. For imaging of live samples, PAAm gels were prepared onto glass bottom dishes (ThermoFisher Scientific).

**Table 1.** Component ratios for the preparation of PAAm hydrogels with different stiffness.

| Hydrogel | AAm |  | BisAAm |  |
| --- | --- | --- | --- | --- |
| | Percentage (%) | Volume (μL) | Percentage (%) | Volume (μL) |
| Soft | 3 | 30 | 0.06 | 50 |
| Medium | 5 | 12 | 0.3 | 60 |
| Rigid | 10 | 325 | 0.3 | 257 |

### ECM protein-functionalization of PAAm hydrogels

PAAm hydrogels prepared on coverslips were transferred to multi-well plates before covering with 0.2 mg/mL sulfosuccinimidyl 6-(4’-azido-2’-nitrophenylamino)hexanoate (sulfo-SANPAH) (Thermo Fisher). Samples were placed in a 365 nm UV light source at a distance of ∼3 inches for 10 min. This step was repeated 3 times. Hydrogels were then washed with 50 mM HEPES buffer (pH 8.5) 3 times before coating with 10 µg/mL ECM protein (fibronectin or laminin-111, as indicated) (Biolamina) in HEPES buffer and overnight incubation at 37°C. Hydrogels were washed with milli-Q water to remove excess protein.

### PAAm hydrogels functionalization quantification

Hydrogels functionalization was quantified by measurement of fibronectin coating intensity. Hydrogels were washed with DPBS, blocked in 2% BSA in DPBS for 1 h at room temperature, and then incubated with mouse monoclonal primary antibody against fibronectin (1:400, Sigma-Aldrich) in blocking solution overnight at 4°C. Hydrogels were then rinsed twice in DPBS/0.1% Triton X-100 and incubated with goat anti-rabbit Cy3-conjugated (1:200, Jackson Immunoresearch) secondary antibody at room temperature for 1 h. Samples were imaged using a Zeiss Observer Z1 epifluorescence inverted microscope. 3 replicates per sample were measured.

### PAAm hydrogels nanoindentation

Nanoindentation measurements were performed using a fiber-optic based nanoindentation device Chiaro (Optics11) mounted on top of an inverted optical microscope Zeiss Axiovert 200M (Zeiss). Measurements were performed following the standardized protocol described by Ciccone *et al.* ^83^ using a cantilever with stiffness (k) 0.51 Nm^-1^ holding spherical tips of radius (R) 3 µm for medium and stiff gels, and 0.028 Nm^-1^ with 28.5 µm for soft gels, respectively. For each experimental condition, at least 3 hydrogels were indented with a minimum of 100 indentations per condition. For each indentation the probe moved at a constant speed of 2 μm/s over a vertical range of 10 μm (displacement control). The forward segment of the collected force-displacement (F-z) curves was analyzed using a custom open-source software ^83^. Curves were first filtered using a Savitzky Golay filter from the SciPy computing stack ^84^ with window length of 25 nm and polynomial order of 3 to remove random noise. After, the point where the probe came into contact with the cell (*z*_0_, *F*_0_) was identified with a thresholding algorithm to convert (F-z) curves into force-indentation (F-δ) curves. To quantify the elastic properties of the gels (Young’s Modulus E), F-δ curves were fitted with the Hertz model (Equation 1) up to a maximum indentation of δ = 0.1 *R*. The Poisson’s ratio (*v*) was taken as 0.5 assuming material’s incompressibility.

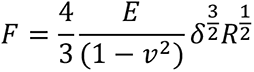

### Cell culture

Murine C2C12 myoblasts (Sigma-Aldrich) were maintained in Dulbecco’s Modified Eagle Medium (DMEM, Invitrogen) with high glucose content, supplemented with 20% Foetal Bovine Serum (FBS, Invitrogen) and 1% antibiotics (P/S) (1 mL of a mixture of 10,000 units/mL of penicillin and 10,000 μg/mL streptomycin per 100 mL of media, ThermoFisher Scientific) in humidified atmosphere at 37°C and 5% CO2.

### Cell viability

Cytotoxicity assay MTT ((3-(4,5-dimethylthiazol-2-yl)-2,5-diphenyltetrazolium bromide) quantitative assay (Promega) was performed to assess cytocompatibility of PAAm hydrogels and borax (Borax España S.A) with C2C12 cells. 1×10^4^ cells/cm^2^ cells were seeded on PAAm hydrogels and metabolic activity was measured after 1, 3, 5 and 7 days of incubation in DMEM supplemented with 20% FBS and 1% P/S. Samples were treated with borax 0.59 or 1.47 mM as required. These concentrations are based in previous works from the group ^28,29^. Cells were then incubated for 2 h with MTT (tetrazolium salt) at 37°C. Formazan was solubilized with DMSO followed by measuring absorbance at 540 nm. 3 biological replicates with 3 technical replicates were measured.

MTT assays were complemented with LIVE/DEAD® viability/cytotoxicity kit (Molecular Probes). Briefly, samples were washed with PBS at 37°C and incubated for 30 min at 37°C with a mixture containing 4 µM ethidium homodimer-1 and 2 µM calcein AM in PBS. Then, the staining solution was removed, and the samples were washed with PBS. Samples were imaged using a ZEISS Axio Observer Z1 epifluorescence inverted microscope. Image processing and analysis was performed using Fiji imaging software. 3 biological replicates with 3 technical replicates were measured.

### Cell proliferation

1×10^4^ cells/cm^2^ cells were seeded on PAAm hydrogels and allowed to adhere for 24 h. Samples were treated with borax 0.59 or 1.47 mM as required. Cells were incubated with alamarBlue^®^ reagent (ThermoFisher Scientific) for 2h protected from the light in humidified atmosphere at 37°C and 5% CO2. Absorbance was read at 570 nm using a Multiskan FC microplate reader (ThermoScientific). 600 nm was used as a reference wavelength. 3 biological replicates with 3 technical replicates were measured.

### Cell adhesion

C2C12 cells were seeded at low density of 5×10^3^ cells/cm^2^ onto functionalized PAAm hydrogels and allowed to adhere for 3 h. Cells were cultured in DMEM with high glucose content, supplemented with 1% P/S and in absence of serum (FBS). After 3 h of culture, cells were washed in DPBS (Gibco) and fixed in 4% formaldehyde solution (Sigma-Aldrich) for 20 min. Samples were treated with borax 0.59 or 1.47 mM as required. 10 cells from 3 different biological replicates were measured.

### Immunostaining

Cells from adhesion and differentiation experiments were rinsed with DPBS and permeabilized with 0.5% Triton x-100 in DPBS at room temperature for 5 min, next blocked in 2% BSA (Sigma-Aldrich) in DPBS for 1 h at room temperature, and then incubated with primary antibodies in blocking solution overnight at 4°C. The samples were then rinsed twice in DPBS/0.1% Triton X-100 and incubated with the secondary antibody and phalloidin (Invitrogen) at room temperature for 1 h. Finally, samples were washed twice in 0.1% Triton X-100 in DPBS before mounting with Vectashield containing DAPI (Vector Laboratories). For cell adhesion studies, mouse monoclonal primary antibody against vinculin (1:400, Sigma-Aldrich), Alexa fluor 488 phalloidin (1:200, Invitrogen) and rabbit anti-mouse Cy3-conjugated (Jackson Immunoresearch, 1:200) secondary antibody were used. For myogenic differentiation studies, mouse monoclonal primary antibody against sarcomeric α-actinin (1:200, Abcam) and rabbit anti-mouse Cy3-conjugated secondary antibody (1:200, Jackson Immunoresearch) were used. For intracellular tension studies, mouse monoclonal primary antibody against phospho-myosin light chain (1:200, Cell Signalling), rabbit monoclonal primary antibody against yes-associated protein 1 (YAP) (1:500, Abcam), Alexa fluor 488 phalloidin (1:200, Invitrogen) and rabbit anti-mouse Cy3-conjugated (1:200, Jackson Immunoresearch) and goat anti-rabbit Cy3-conjugated (1:200, Jackson Immunoresearch) secondary antibodies were used.

Samples were imaged using a Zeiss Observer Z1 epifluorescence inverted microscope or Zeiss LSM900 confocal microscope using Micro-Manager or ZEN software, respectively. Image processing and analysis was performed using Fiji imaging software.

### Actin flow

C2C12 cells were transfected using the Neon transfection system (ThermoFisher Scientific) following manufacturer’s protocol. Plasmid used was LifeAct-GFP (Ibidi). Parameters used to achieve cell transfection were 1650 V, 10 ms, 3 pulses, with 5 μg of DNA. Transfected cells were cultured for 24 h.

Cells were seeded at 1×10^4^ cells/cm^2^ on PAAm hydrogels and allowed to adhere for 24 h. Cells were imaged using a ZEISS LSM900 confocal microscope with a 40x oil-immersion objective and ZEN software. Images were taken for 4 min at 1 frame every 2 seconds at 488 nm. Actin flow was determined by kymographs in the Fiji software. Samples were treated with borax 0.59 or 1.47 mM as required. 5 cells with at least 5 different flow areas per cell were measured.

### Cells nanoindentation

Nanoindentation measurements were performed using a fiber-optic based nanoindentation device Chiaro (Optics11) mounted on top of an inverted optical microscope Zeiss Axiovert 200M (Zeiss). Measurements were performed following the standardized protocol described by Ciccone *et al*. ^83^ using cantilevers with stiffness (k) of 0.020 or 0.022 Nm^-1^ holding spherical tips of radius (R) 3.5 μm and 3 μm, respectively.

C2C12 myoblasts were plated on functionalized glasses at a density of 1×10^4^ cells/cm^2^ and allowed to adhere for 24 h. All measurements were performed at 37°C using an on-stage Incubator (Okolab) in standard culture medium.

For each experimental condition, at least 10 cells were indented by performing 9 repeated indentations on each single cell, with subsequent indentations being spaced 1 μm apart. For each indentation the probe moved at a speed of 2 μm/s over a vertical range of 10 μm.

The forward segment of the collected force-displacement (F-z) curves was analyzed using a custom open-source software ^83^. Curves were first filtered using a Savitzky Golay filter from the SciPy computing stack ^84^ with window length of 25 nm and polynomial order of 3 to remove random noise. After, the point where the probe came into contact with the cell (*z*_0_, *F*_0_) was identified with a thresholding algorithm to convert (F-z) curves into force-indentation (F-δ) curves. To quantify the elastic properties of the gels (Young’s Modulus E), F-δ curves were fitted with the Hertz model (Equation 1) up to a maximum indentation of δ = 0.1 *R*. The Poisson’s ratio (*v*) was taken as 0.5 assuming material’s incompressibility.

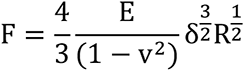

### Brillouin microscopy

Brillouin microscopy (LifeMachinery) was performed by using a 20x objective to illuminate the sample with a 660 nm laser. The backscattered light was coupled into a single-mode fiber and analyzed using a VIPA cross-axis spectrometer. Each pixel in the Brillouin maps came from one Brillouin spectrum. In the VIPA spectrometer, the frequencies of the light were separated in space and imaged on a high-sensitivity Orca Fusion camera. Both the stoke and anti-stoke peaks, which correspond to lower and higher energy (compared to Rayleigh) scattered radiation, were fitted with a Lorentzian function. The frequency (Brillouin) shift, measured as the displacement of the Brillouin peaks with respect to the Rayleigh, was taken from the LabView software (Light Machinery). Before the measurements, a first calibration with an acrylic cube and Spectralock software was performed. To minimise the laser signal, the etalon pressure was modified by starting with increments of 100 and then 10. Once the laser peak was minimized, a stripe overlay was performed to ensure that the spectra were well extracted with the optimal Brillouin signal. Finally, the collimator coupling was optimized to increase the signal of the Brillouin peaks. A second calibration was then performed with the sample to locate the laser blobs and a striped overlay was then performed. After this, the background was subtracted by closing the laser shutter and making an average of 10 spectra. The laser shutter was then opened and the Brillouin peaks were selected in the LabView software. To take the maps of the sample, a square was placed in the camera window and the size of the square and the size of the step size between measurements was selected. 10 cells from 3 different biological replicates were measured.

### Traction Force Microscopy (TFM)

Carboxylate-modified 0.2 µm FluoSpheres (Life Technologies) were prepared by sonicating the stock for 10 min, then diluted 1:30 in milli-Q water and further sonicated for 15 min. Immediately after sonication, 1:20 FluoSpheres were incorporated into the PAAm hydrogels on coverslips before functionalization, as previously described.

Cells were seeded at 1×10^4^ cells/cm^2^ on PAAm hydrogels and allowed to adhere for 24 h. Using an EVOS FL Auto microscope (Life Technologies) with the incubator set at 37°C and 5% CO2 at 20x magnification, Z-stack images were taken through the cells (brightfield channel) and FluoSpheres embedded in the hydrogels (Texas Red channel) before and after cell trypsinization. Cell traction forces were determined using ImageJ software by tracking the displacement of the FluoSpheres and then reconstructing the force field from the displacement data using the iterative particle image velocimetry (PIV) and Fourier transform traction cytometry (FTTC) plugins ^85^ in ImageJ software, respectively. The stress maps obtained were modified in ParaView software to plot more accurate scales. At least 30 cells from 10 different beacons per hydrogel were analyzed.

### Gene expression

Total RNA was extracted from C2C12 cultured for 4, 8, 24 or 96 h under different experimental conditions using RNeasy^®^ Micro Kit (Qiagen). RNA quantity and integrity was measured with a NanoDrop 1000 (ThermoScientific). Then 500 ng of RNA were reverse transcribed using the QuantiTect^®^ Reverse Transcription Kit (Qiagen). Real-time qPCR was performed using Quantinova SYBR^®^ Green PCR kit (Qiagen) and 7500 Real Time PCR system (Applied Biosystems). The reactions were run in triplicate for both technical and biological replicas. The primers used for amplification were designed from sequences found in the GenBank database and included: *NaBC1* (Gene ID: 269356; Fw: 5′-GAGGTTCGCTTTGTCATCCTGG-3′, Rev: 5′-ATGCCAGTGAGCTTCCCGTTCAG-3′), *AKT* (Gene ID: 11651; Fw: 5′-GGACTACTTGCACTCCGAGAAG-3′, Rev: 5′-CATAGTGGCACCGTCCTTGATC-3′), *mTOR* (Gene ID: 56717; Fw: 5′-AGAAGGGTCTCCAAGGACGACT-3′, Rev: 5′-GCAGGACACAAAGGCAGCATTG-3′), *GDF11* (Gene ID: 14561; Fw: 5′-TTTCGCCAGCCACAGAGCAACT-3′, Rev: 5′-CTCTAGGACTCGAAGCTCCATG-3′), *MyoD* (Gene ID: 17927; Fw: 5′-GCACTACAGTGGCGACTCAGAT-3′, Rev: 5′-TAGTAGGCGGTGTCGTAGCCAT-3′), *MYOGENIN* (Gene ID: 17928; Fw: 5′-CCATCCAGTACATTGAGCGCCT-3′, Rev: 5′-CTGTGGGAGTTGCATTCACTGG-3′), *VEGFR* (Gene ID: 14254; Fw: 5′-TGGATGAGCAGTGTGAACGGCT-3′, Rev: 5′-GCCAAATGCAGAGGCTTGAACG-3′), *INSR* (Gene ID: 16337; Fw: 5′-AGATGAGAGGTGCAGTGTGGCT-3′, Rev: 5′-GGTTCCTTTGGCTCTTGCCACA-3′), *IL-GFR* (Gene ID: 16001; Fw: 5′-CGGGATCTCATCAGCTTCACAG-3′, Rev: 5′-TCCTTGTTCGGAGGCAGGTCTA-3′), *INTEGRIN ALPHA v* (Gene ID: 16410; Fw: 5′-GTGTGAGGAACTGGTCGCCTAT-3′, Rev: 5′-CCGTTCTCTGGTCCAACCGATA-3′), *INTEGRIN ALPHA 5* (Gene ID: 16402; Fw: 5′-ACCTGGACCAAGACGGCTACAA-3′, Rev: 5′-CTGGGAAGGTTTAGTGCTCAGTC-3′), *INTEGRIN ALPHA 7* (Gene ID: 16404; Fw: 5′-TCTGTCAGAGCAACCTCCAGCT-3′, Rev: 5′-CTATGAACGGCTGCCCACTCAA-3′), *INTEGRIN BETA 1* (Gene ID: 16412, Fw: 5′-CTCCAGAAGGTGGCTTTGATGC-3′, Rev: 5′-GTGAAACCCAGCATCCGTGGAA-3′), *INTEGRIN BETA 3* (Gene ID: 16416, Fw: 5′-GTGAGTGCGATGACTTCTCCTG-3′, Rev: 5′-CAGGTGTCAGTGCGTGTAGTAC-3′), *INTEGRIN BETA 5* (Gene ID: 16419, Fw: 5′-TTTCGCCAGCCACAGAGCAACT-3′, Rev: 5′-CTCTAGGACTCGAAGCTCCATG-3′). *GAPDH* (Gene ID: 14433; Fw: 5′-CATCACTGCCACCCAGAAGACTG-3′, Rev: 5′-ATGCCAGTGAGCTTCCCGTTCAG-3′) was used as a housekeeping gene. Ct value was used for quantification by the comparative Ct method. Sample values were normalized to the threshold value of housekeeping gene GAPDH: ΔCT = CT(gene of interest) − CT(GAPDH). The Ct value of the control (cell culture plate) was used as a reference. ΔΔCT = ΔCT(experiment) − ΔCT(control). mRNA expression was calculated by the following equation: fold change = 2^-ΔΔCT^. 3 biological replicates with 3 technical replicates were measured.

### NaBC1-integrins colocalization

C2C12 myoblasts were plated on PAAm hydrogels at a density of 1×10^4^ cells/cm^2^ and allowed to adhere for 24 h. Samples were treated with borax 0.59 or 1.47 mM as required. Colocalization of NaBC1/Integrin α_v_, NaBC1/α_5_ and NaBC1/β_4_ experiments were performed using DUOLINK^®^ PLA system (Sigma-Aldrich) following the manufacturer’s instructions. Specific primary antibodies used were: anti-NaBC1 (Invitrogen, 1:200), anti-integrin α_v_ (Abcam, 1:500), anti-integrin α_5_ (Abcam, 1:500) and anti-integrin β_4_ (Abcam, 1:500). For image quantification of colocalization fluorescent dots, at least 30 individual cells were imaged for each condition using a ZEISS LSM900 confocal microscope.

### Cell senescence

Cell senescence was measured with the CellEvent^™^ Senescence Green Detection Kit (Invitrogen). The kit is based on the CellEvent^™^ Senescence Green Probe, which is a fluorescent reagent containing two galactoside moieties, making it specific to the typical senescence marker β-galactosidase. The enzyme-cleaved product is retained within the cell due to covalent binding of intracellular proteins and emits a fluorogenic signal that has excitation/emission maxima of 490/514 nm. C2C12 cells were seeded at 1×10^4^ cells/cm^2^ on PAAM hydrogels and allowed to adhere for 24 h. Samples were treated with borax 0.59 or 1.47 mM as required. Cells were washed with DPBS (Gibco) and fixed in 4% formaldehyde solution (Sigma-Aldrich) for 20 min. Samples were washed within BSA 1% in DPBS and incubated in the dark with CellEvent^™^ Senescence Green Probe for 2 hours at 37°C without CO2. After incubation, samples were washed three times with DPBS, and fluorescence was quantified in a Multiskan FC microplate reader (ThermoScientific) using an Alexa Fluor^™^ 488/FITC filter set. Samples were imaged with a Zeiss LSM900 confocal microscope. 3 biological replicates with 3 technical replicates were measured.

### Total ATP content

C2C12 cells were seeded at 1×10^4^ cells/cm^2^ on PAAM hydrogels. After 24 h, cells were washed twice with PBS and lysed using 0.5% Triton-X100 in PBS. After centrifugation at 11,000 × g for 3 min, 5 µl of total lysate were used in triplicates for assessment of total ATP content using the ATP Determination Kit (ThermoFisher Scientific) following the manufacturer’s instructions. Luminescence was monitored at 560 nm using a Multiskan FC microplate reader (ThermoScientific). 3 biological replicates with 3 technical replicates were measured.

### Mitochondrial ATP content

Mitochondrial ATP content was assessed with the BioTracker^TM^ ATP-Red dye (Millipore), a live cell red fluorescent imaging probe for ATP. The probe specifically reports ATP content in the mitochondrial matrix of living cells. The probe without ATP forms a closed ring structure that is not fluorescent. In the presence of the negatively charged ATP, the covalent bonds between boron and ribose in the probe are broken and the ring opens, causing the probe to be fluorescent.

C2C12 cells were seeded at 1×10^4^ cells/cm^2^ on PAAM hydrogels. After 24 h, cells were incubated for 1 h with 200 nM MTG. Then, cells were washed twice with PBS and incubated for 15 min with 5 µM BioTracker^TM^ ATP-Red dye in medium at 37°C and 5% CO2. Then, the cells were washed twice with medium and fresh medium was added. Imaging was performed using a ZEISS LSM900 confocal microscope. Analysis was performed with ZEN software by measuring the average ATP red fluorescence intensity inside a region of interest generated by the MTG area. At least 10 cells from 3 different biological replicates were measured.

### Chemical procedure for the preparation of Fluorescein-labelled boronic acid

Chemical synthesis reactions were performed employing commercial reagents and solvents without additional purification unless otherwise noted. Solvents for synthesis were purchased from Scharlab while chemicals were purchased from usual commercial sources. Particularly, 4-(Aminomethyl)phenylboronic acid pinacol ester hydrochloride, HCl in methanol (1.25M), triethylamine and sodium metaperiodate were purchased from Sigma-Aldrich while Fluorescein isothiocyanate (FITC, 95% mixture of isomers) was purchased from ABCR. Analytical thin layer chromatography (TLC) was performed on precoated aluminium silica gel sheets. NMR spectra were recorded in a Bruker AV250 equipment employing deuterated solvents purchased from Sigma-Aldrich. Chemical shifts were reported relative to the remainder ^1^H of solvents. Coupling constants J were given in Hz. Resonance patterns were designated with the notations s (singlet), d (doublet), m (multiplet), br s (broad singlet). FTIR spectra were recorded using a Nexus (ThermoFisher Scientific) spectrometer equipped with a Smart Golden Gate ATR accessory. Mass spectrometry spectra were recorded in a Bruker HTC ion trap mass spectrometer.

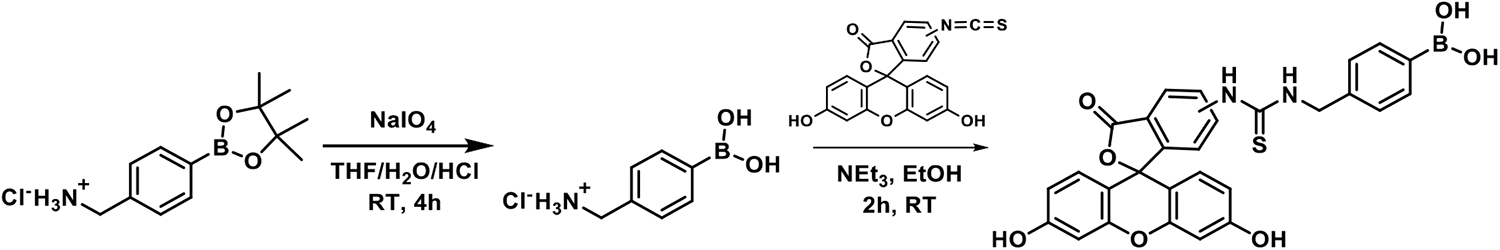

Title compound, 4-(fluoresceinyl)thioureidomethylphenylboronic acid was prepared in a two-step procedure from 4-(aminomethyl)phenylboronic acid pinacol ester. First, the cleavage of the pinacol ester was achieved upon oxidative cleavage of the pinacol C-C bond. Briefly, NaIO_4_ (240 mg, 1.1 mmol 1.2 equiv.) and the ammonium-containing boronic ester (250 mg, 0.93 mmol, 1.0 equiv.) were dissolved in a mixture of THF (8 mL), water (2 mL) and hydrochloric acid (0.1 mL). This mixture was stirred at room temperature for 4h. To isolate boronic acid, solvents were removed under vacuum and the crude dissolved in ethanol. Inorganic salts were filtered off and the crude was then treated with a solution of hydrogen chloride in methanol (5mL) to form the hydrochloride. Finally, intermediate compound was isolated after solvent removal and precipitation with a mixture of chloroform and heptane. The product was isolated as a white solid (145 mg, 83%) and employed for the next step without further purification. ^1^H NMR (250 MHz, MeOD) δ 7.70 (d, J = 7.2 Hz, ^1^H), 7.45 (d, J = 7.4 Hz, ^1^H), 4.12 (s, ^1^H). MS(ESI^+^), m/z calculated for (M+H)^+^ [C_7_H_11_BNO_2_]^+^: 152.09, found: 152.1. Melting point: >180°C (decomposition).

The preparation of title compound was performed by direct reaction of the as prepared aminomethyl phenylboronic acid hydrochloride (9.4 mg, 0.05 mmol) with stoichiometric amounts of FITC (19.5 mg, 0.05 mmol) and a stock solution of triethylamine (0.01 M) in EtOH (5 mL). The reaction was stirred for 2 h in the dark, evaporated and filtered through a flash chromatography column employing silica gel and EtOAc as eluent. Final compound was obtained as an orange solid (26 mg, 96%). ^1^H NMR (250 MHz, MeOD) δ 8.05 (d, J = 1.8 Hz, ^1^H), 7.75 (dd, J = 8.2, 2.0 Hz, ^1^H), 7.62 (br. d, J = 7.2 Hz, ^2^H), 7.38 (d, J = 7.9 Hz, ^2^H), 7.21 (d, J = 8.5 Hz, ^1^H), 7.07-6.98 (m, ^3^H), 6.74 – 6.51 (m, ^7^H), 4.88 (s, ^2^H). MS(ESI^+^), m/z calculated for (M+H)^+^ [C_28_H_21_BN_2_O_7_S]^+^: 540.12, found: 540.1.

### Subcellular localization

C2C12 myoblasts were plated on PAAm hydrogels at a density of 1×10^4^ cells/cm^2^ and allowed to adhere for 24 h. Cells were treated with FITC-B for 1 h at 37°C. After washing with DPBS, cells were incubated with 75 nM LysoTracker^®^ Red DND-99 (Invitrogen), 100 nM MitoTracker^®^ Red CMXRos (Invitrogen) or 1 µM ER-tracker^®^ Red (Invitrogen) for 1 h at 37°C. Samples were imaged with a Zeiss LSM980 confocal microscope and an incubator to maintain the conditions constant at 37°C and 5% CO2. Manders overlapping coefficient (MOC) was determined with ZEN software. 20 cells from 3 different biological replicates were measured.

### Fluorescence recovery after photobleaching (FRAP)

C2C12 myoblasts were plated on PAAm hydrogels at a density of 1×10^4^ cells/cm^2^ and allowed to adhere for 24 h. Samples were treated with FITC-B for 1 h, as required. Then, samples were washed with DPBS three times and imaged with a Zeiss LSM980 confocal microscope. 488 laser was used to bleach 100% of fluorescence in the indicated areas (nucleus, cytoplasm, mitochondria, lysosomes, endoplasmic reticulum and focal adhesions) for 2 ms. Fluorescence recovery was measured for 4 minutes after bleaching. 10 cells from 3 different biological replicates were measured.

### Myogenic differentiation

C2C12 cells were plated on PAAm hydrogels at high seeding density of 2×10^4^ cells/cm^2^ in differentiation medium for myotube formation (DMEM high glucose content supplemented with 1% Insulin-Transferrin-Selenium (ITS, Gibco) and 1% P/S. Samples were treated with borax 0.59 or 1.47 mM as required. Differentiation medium was changed every 2 days. After 4 days of culture, cells were washed in DPBS (Gibco) and fixed in 4% formaldehyde solution (Sigma-Aldrich) for 20 min. 3 biological replicates with 3 technical replicates were measured.

### NaBC1 silencing

C2C12 cells were seeded at 6×10^4^ cells/cm^2^ on PAAm hydrogels in DMEM with high glucose content, supplemented with 20% FBS and 1% P/S in humidified atmosphere at 37°C and 5% CO2. After 24 h, cells were washed with Opti-MEM reduced serum medium (ThermoFisher Scientific) and transfected using pre-designed MISSION^®^ esiRNA (Sigma-Aldrich) against mouse NaBC1 in X-tremeGENE siRNA Transfection Reagent (Roche), following manufacturer’s instructions. MISSION^®^ esiRNA Fluorescent Universal Negative Control 1 Cy3 (NC, Sigma-Aldrich) was used as a control of transfection efficiency. NaBC1 silencing was corroborated by evaluation of NaBC1 mRNA expression levels.

### VD-1 transfection

C2C12 cells were transfected using the Neon transfection system (ThermoFisher Scientific) following manufacturer’s protocol. Plasmid used was VD-1-GFP (kindly gifted by Pere Roca-Cusachs). Parameters used to achieve cell transfection were 1650 V, 2 ms, 3 pulses, with 5 μg of DNA. Transfected cells were cultured for 24 h. Following VD-1 transfection, cell morphology, immunostaining, and actin flow assays were performed, as previously described.

### Image analysis

For PAAm hydrogels functionalization measurement, staining intensity of immunofluorescence images (fibronectin) was quantified by Fiji software.

For focal adhesions analysis, vinculin images were segmented by ImageJ, using Trainable Weka Segmentation plugin to create a binary mask. After segmentation, focal adhesion number, length and area were determined using different commands of the same software. Cell morphology was analyzed by calculation of different parameters using Fiji software.

For intracellular tension studies, staining intensity of immunofluorescence images (phospho-myosin light chain) was quantified by Fiji software.

YAP expression was plotted as a nuclear/cytoplasmic ratio; this was performed by measuring nuclear and cytoplasmic YAP expression independently and calculated as follows:

First, cytoplasmic area was defined, where A*cell* is the area of the entire cell and A*nuc* is the area of cell nucleus.

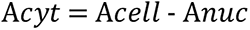

Second, YAP’s integrated density fluorescence in the cytoplasm was calculated, where YAP*cell* is the integrated density of YAP in the entire cell and YAP*nuc* is the integrated density of YAP in the cell nucleus.

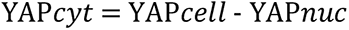

Finally, YAP’s integrated density fluorescence nucleus/cytoplasm ratio is calculated, where YAP*nuc* is the integrated density of YAP in the nucleus, A*nuc* is the area of the nucleus, YAP*cyt* is the integrated density of YAP in the cytoplasm, and A*cyt* is the area of the cell cytoplasm.

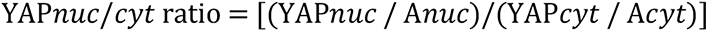

For myogenic differentiation analysis, total nuclei per image were counted from myotube images from differentiation experiments using the particle analysis command in Fiji software. The segmented DAPI channel image was subtracted from the Cy3 channel segmented image, and the remaining nuclei were counted and assigned to non-differentiated cells. The fusion index expressed in %, was calculated subtracting the non-differentiated nuclei from the total nuclei counted. Myotubes were only considered when 3 or more nuclei were aligned inside cells.

## Statistical analysis

Data were analyzed using GraphPad Prism software where normality tests were performed to determine whether to select parametric or non-parametric tests. Then appropriate one-way ANOVA, two-way ANOVA or t-tests, for multiple or pairwise comparisons respectively, were used. Statistical differences were defined by p values and confidence intervals were indicated with a *; * ≤ 0.05, ** ≤ 0.01, *** ≤ 0.001, **** ≤ 0.0001.

## Acknowledgments

M.S-S is grateful for financial support from the European Research Council AdG (Devise, 101054728) and EPSRC HT2050 grant (EP/X033554/1). P.R acknowledges support by grant PID2021-126012OB-I00 funded by MCIN/AEI/10.13039/501100011033 and by ERDF a way of making Europe, and by CIBER (CB06/01/1026). J.G-V acknowledges the funding from the European Union-NextGenerationEU program. The authors acknowledge Pere Roca-Cusachs for the VD-1-GFP and LifeAct-RFP plasmids. IBEC is member of CERCA Programme / Generalitat de Catalunya.

## Author contributions

Conceptualization, P.R.T. and M.S.S.; Methodology, J.G.V, G.C. and E.B.E.; Investigation, J.G.V, G.C., E.B.E. and A.R.N.; Writing – Original Draft, J.G.V.; Writing –Review & Editing, J.G.V, G.C., P.R.T. and M.S-S.; Visualization, J.G.V.; Funding Acquisition, P.R.T. and M.S-S.; Resources, R.C., P.R.T. and M.S-S.; Supervision, P.R.T. and M.S-S.

## Supplemental information

Figures S1-S21.

## Data statement

The data reported in this paper are available from the lead contact upon request.

