## Supplemental figures for "NaBC1 boron transporter enables myoblast response to substrate rigidity via fibronectin-binding integrins"

### Supplemental Information

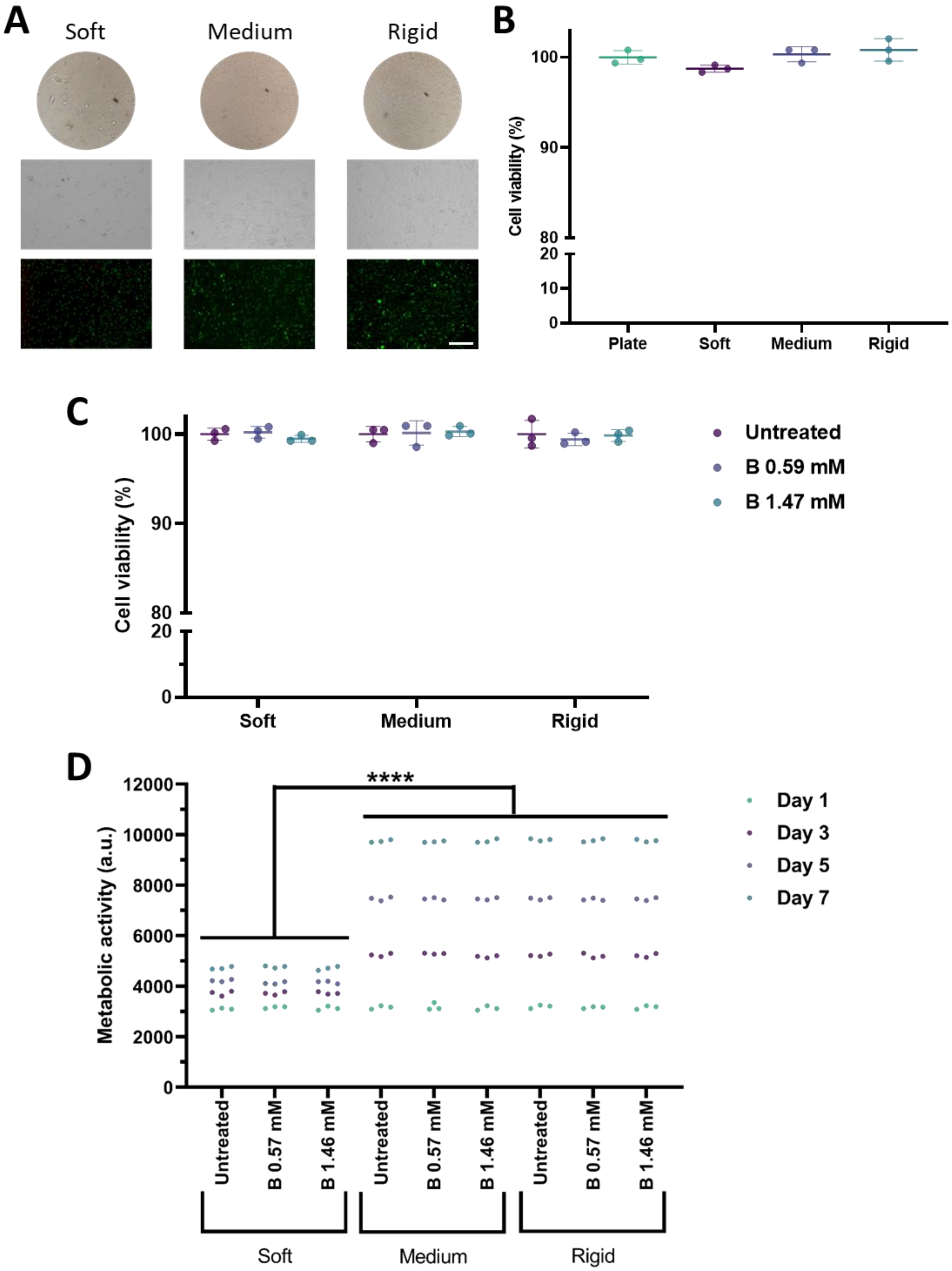

**Figure S1. PAAm hydrogels present excellent biocompatibility.** A: Representative images of C2C12 myoblasts seeded on PAAm hydrogels with different stiffness functionalised with fibronectin. Red: ethidium homodimer-1; Green: calcein AM. Scale bar: 100 µm. B: Quantification of cell viability of C2C12 myoblasts seeded on PAAm hydrogels with different stiffness functionalised with fibronectin. *n*: 3 biological replicates with 3 technical replicates. C: Quantification of cell viability of C2C12 myoblasts seeded on PAAm hydrogels with different stiffness functionalised with fibronectin and stimulated with soluble boron (0.59 and 1.47 mM). *n*: 3 biological replicates with 3 technical replicates. D: Quantification of cell proliferation of C2C12 myoblasts seeded on PAAm hydrogels with different stiffness functionalised with fibronectin and stimulated with soluble boron (0.59 and 1.47 mM) for up to 7 days. *n*: 3 biological replicates with 3 technical replicates. Data are represented as Mean ± Standard Deviation, and differences are considered significant for p ≤ 0.05 using one-way ANOVAs (Tukey's multiple comparisons tests) for multiple comparisons. ****p ≤ 0.0001

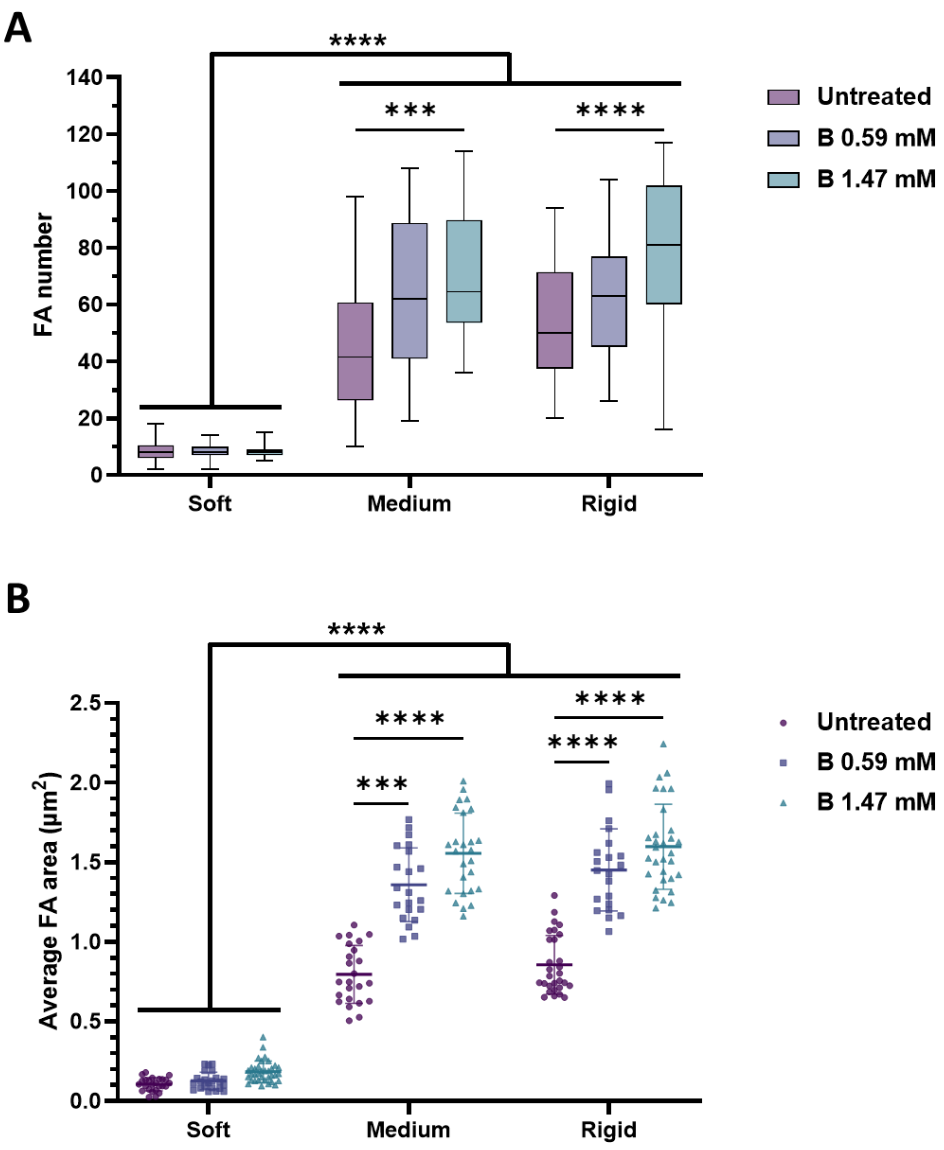


**Figure S2. NaBC1 stimulation triggers the formation of FA in myoblasts on fibronectin.** Quantification of the number (A) and average area (B) of focal adhesions in C2C12 myoblasts seeded on PAAm hydrogels with different stiffness functionalised with fibronectin and stimulated with soluble boron (0.59 and 1.47 mM). *n* = 10 cells from 3 different biological replicates. Data are represented as Mean ± Standard Deviation, and differences are considered significant for p ≤ 0.05 using two-way ANOVAs (Tukey's multiple comparisons tests) for multiple comparisons. ***p ≤ 0.001, ****p ≤ 0.0001

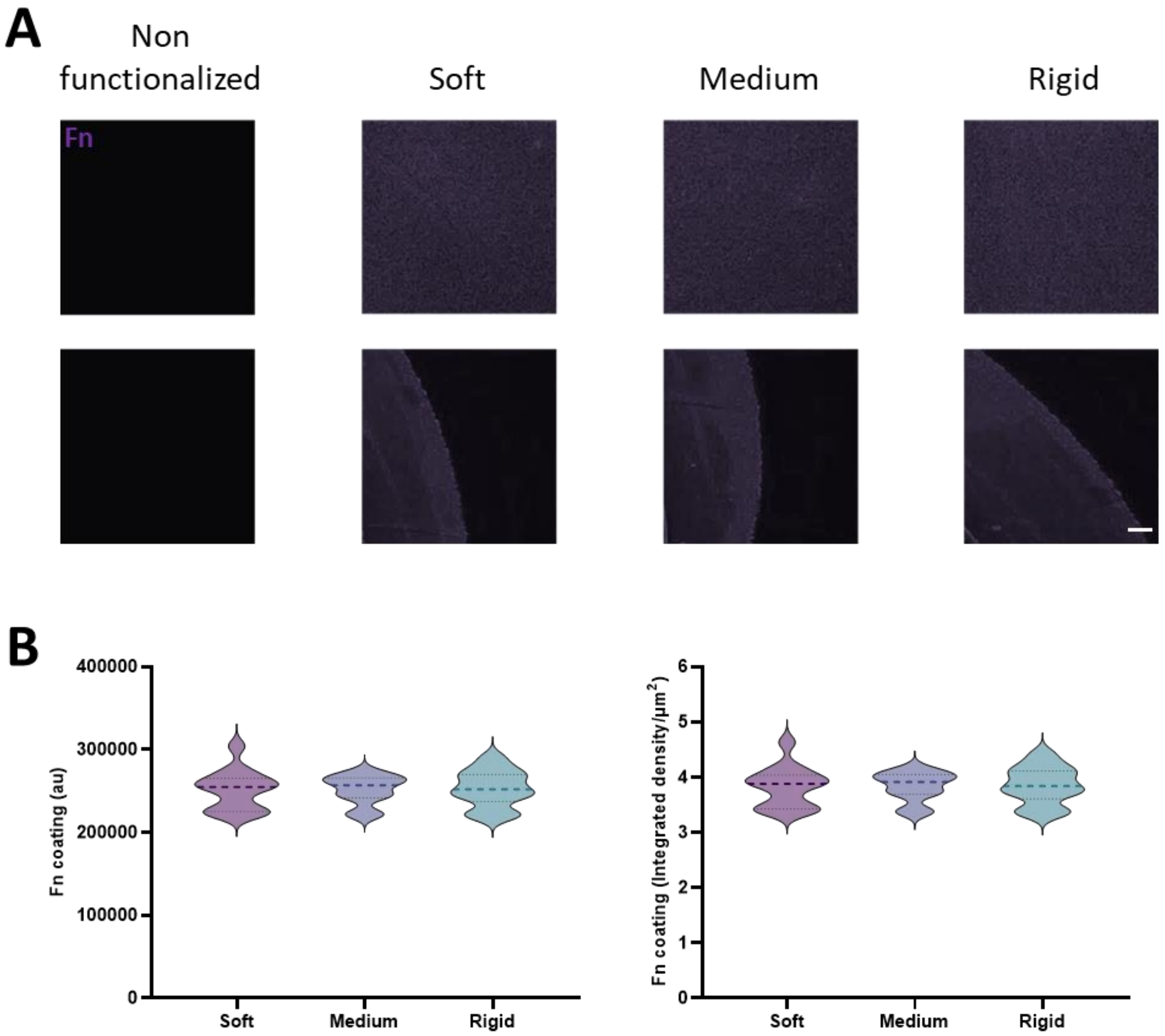


**Figure S3. Fibronectin functionalisation of PAAM hydrogels is homogeneous in all rigidities.** A: Representative images of PAAm hydrogels with different stiffness functionalised with fibronectin. Magenta: fibronectin. Scale bar: 200 µm. B: Quantification of fibronectin coating of PAAm hydrogels with different stiffness. *n*: 3 biological replicates with 3 technical replicates. Data are represented as Mean ± Standard Deviation, and differences are considered significant for $p \leq 0.05$ using one-way ANOVAs (Tukey's multiple comparisons tests) for multiple comparisons.

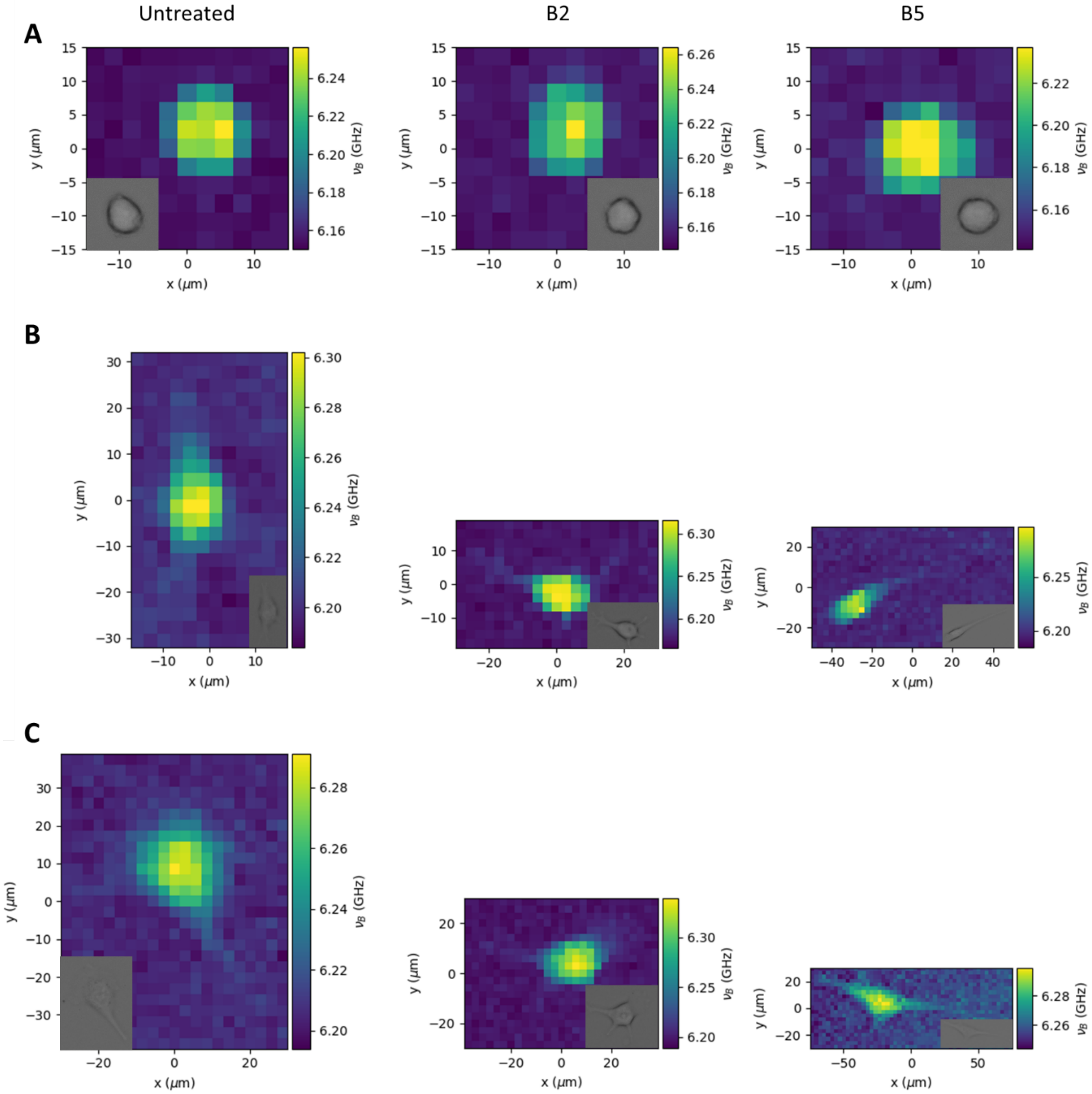


**Figure S4. NaBC1 increases cell stiffness on fibronectin in dependence of substrate rigidity.** Representative Brillouin maps of C2C12 myoblasts seeded on PAAm hydrogels with different stiffness functionalised with fibronectin and stimulated with soluble boron ions (0.59 and 1.47 mM).

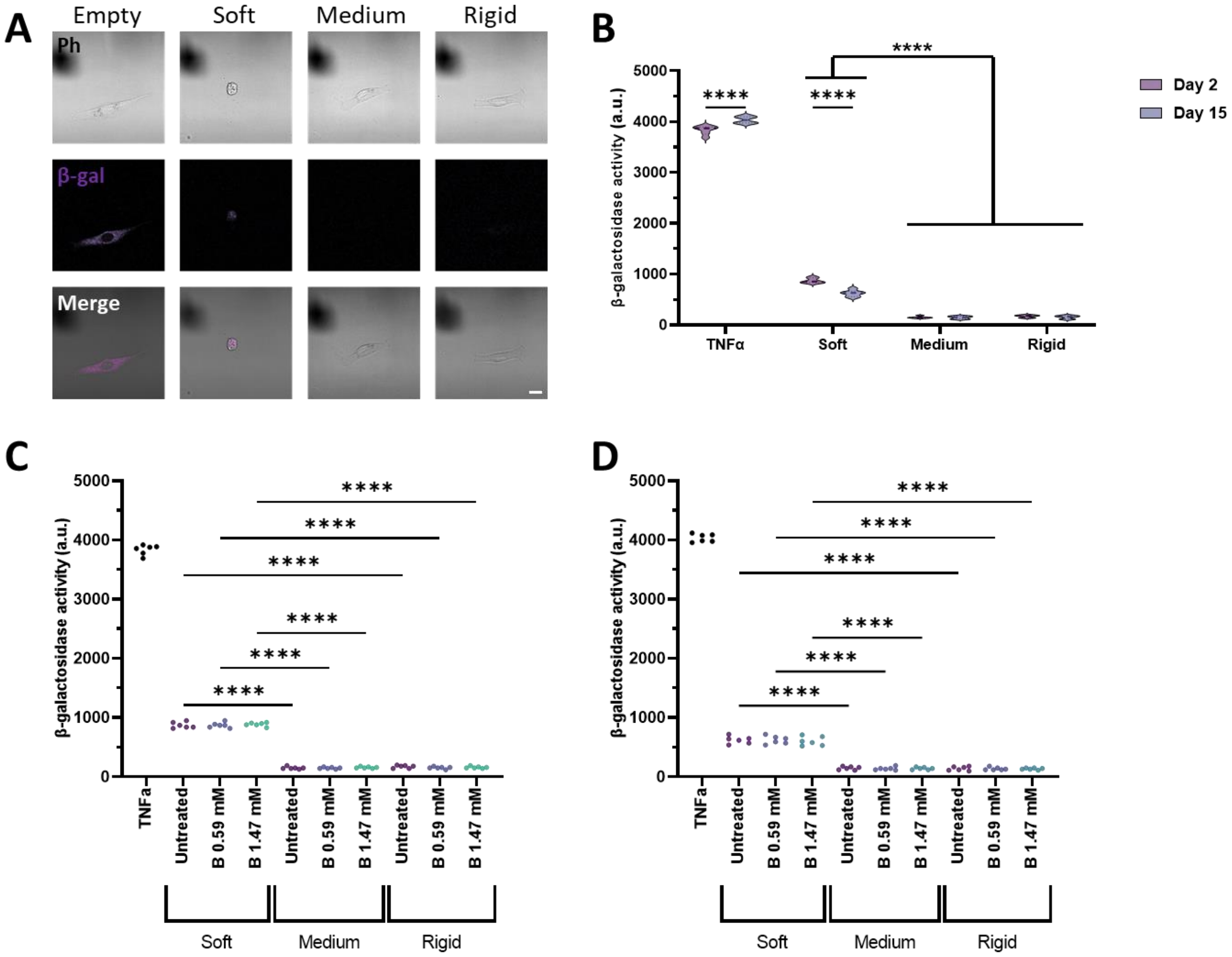


**Figure S5. Cells undergo senescence on soft substrates and it is not reverted by NaBC1 stimulation.** A: Representative images of C2C12 myoblasts seeded on PAAm hydrogels with different stiffness functionalised with fibronectin. Grey: Phase contrast. Magenta: β-galactosidase. fibronectin. Scale bar: 20 µm B: Quantification of β-galactosidase activity in C2C12 myoblasts seeded on PAAm hydrogels with different stiffness functionalised with fibronectin for up to 15 days. *n*: 3 biological replicates with 3 technical replicates. C: Quantification of β-galactosidase activity in C2C12 myoblasts seeded on PAAm hydrogels with different stiffness functionalised with fibronectin and stimulated with soluble boron ions (0.59 and 1.47 mM). *n*: 3 biological replicates with 3 technical replicates. Data are represented as Mean ± Standard Deviation, and differences are considered significant for $p \leq 0.05$ using one-way ANOVAs or two-way ANOVAs (Tukey's multiple comparisons tests) for multiple comparisons. ****$p \leq 0.0001$

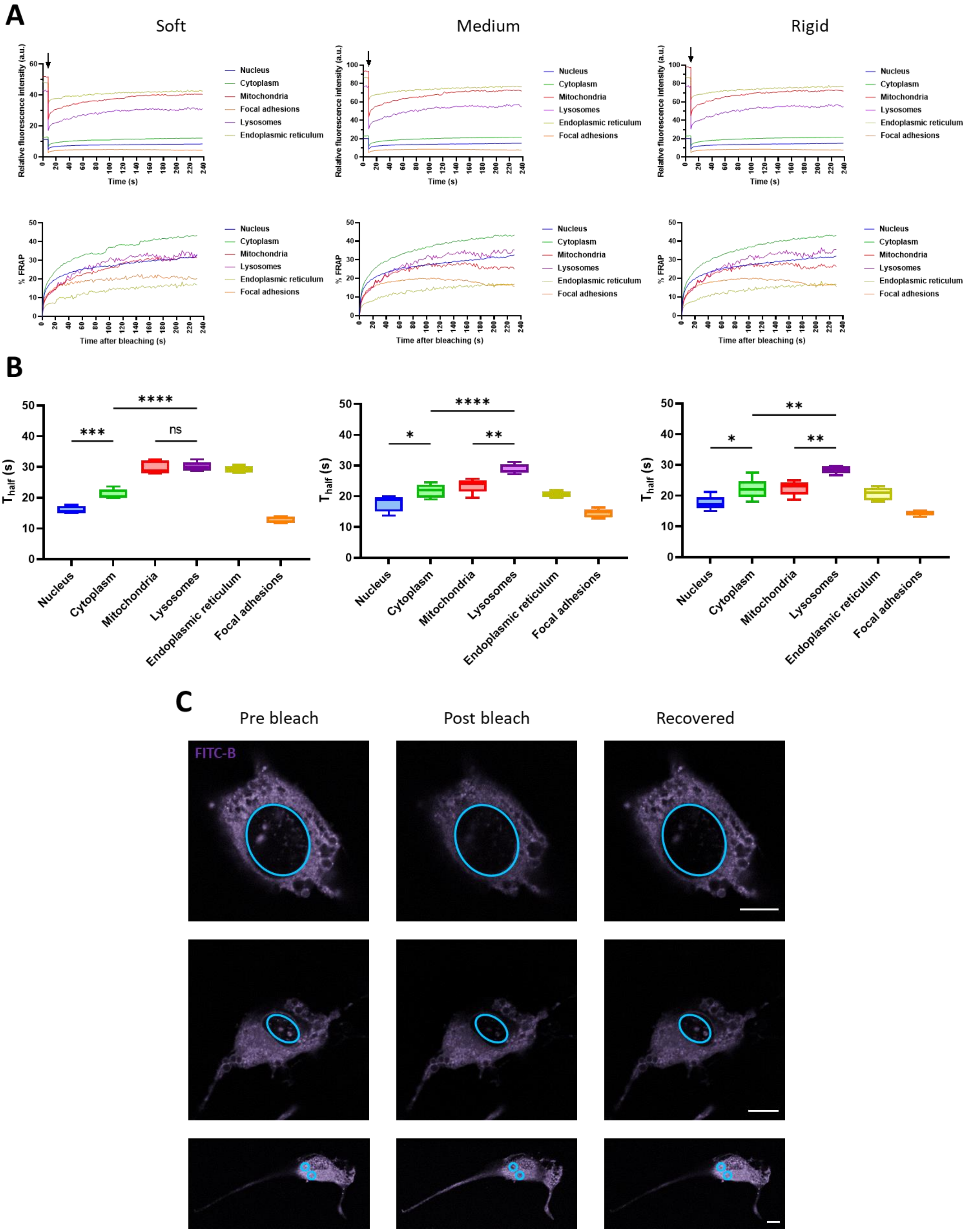


**Figure S6. B subcellular localisation and dynamics.** A: Above: profiles of FITC-labelled B in C2C12 myoblasts seeded on PAAm hydrogels with different stiffness functionalised with fibronectin stimulated with soluble boron. Arrows indicate bleaching time. Below: percentage of signal recovered after bleaching. B: Quantification of half-life of FITC-labelled B in C2C12 myoblasts. *n* = 10 cells from 3 different biological replicates. C: Representative images of C2C12 myoblasts seeded on PAAm hydrogels with different stiffness functionalised with fibronectin stimulated with soluble boron ions. Magenta: FITC-labelled B; Cyan circles: bleached area. Scale bars: 20 µm. Data are represented as Mean ± Standard Deviation, and differences are considered significant for $p \leq 0.05$ using one-way ANOVAs (Tukey's multiple comparisons tests) for multiple comparisons. *p ≤ 0.05, **p ≤ 0.01, ***p ≤ 0.001, ****p ≤ 0.0001

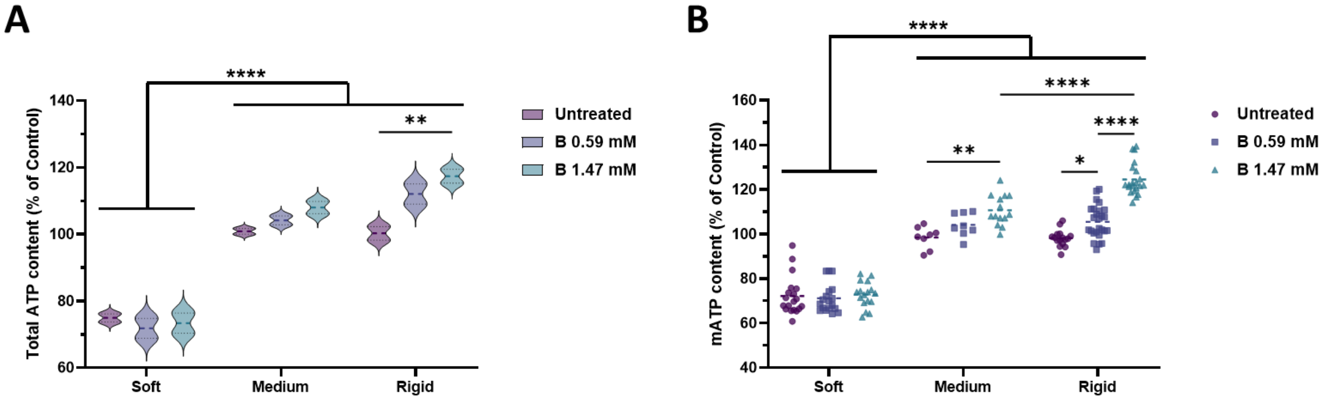


**Figure S7. NaBC1 increases total and mitochondrial ATP content on fibronectin in dependence of substrate stiffness.** A: Quantification of total content of C2C12 myoblasts seeded on PAAm hydrogels with different stiffness functionalised with fibronectin and stimulated with soluble boron (0.59 and 1.47 mM). *n*: 3 biological replicates with 3 technical replicates. B: Quantification of mitochondrial content of C2C12 myoblasts seeded on PAAm hydrogels with different stiffness functionalised with fibronectin and stimulated with soluble boron (0.59 and 1.47 mM). *n*: at least 10 cells from 3 biological replicates. Data are represented as Mean ± Standard Deviation, and differences are considered significant for $p \leq 0.05$ using two-way ANOVAs (Tukey's multiple comparisons tests) for multiple comparisons. *p ≤ 0.05, **p ≤ 0.01, ****p ≤ 0.0001

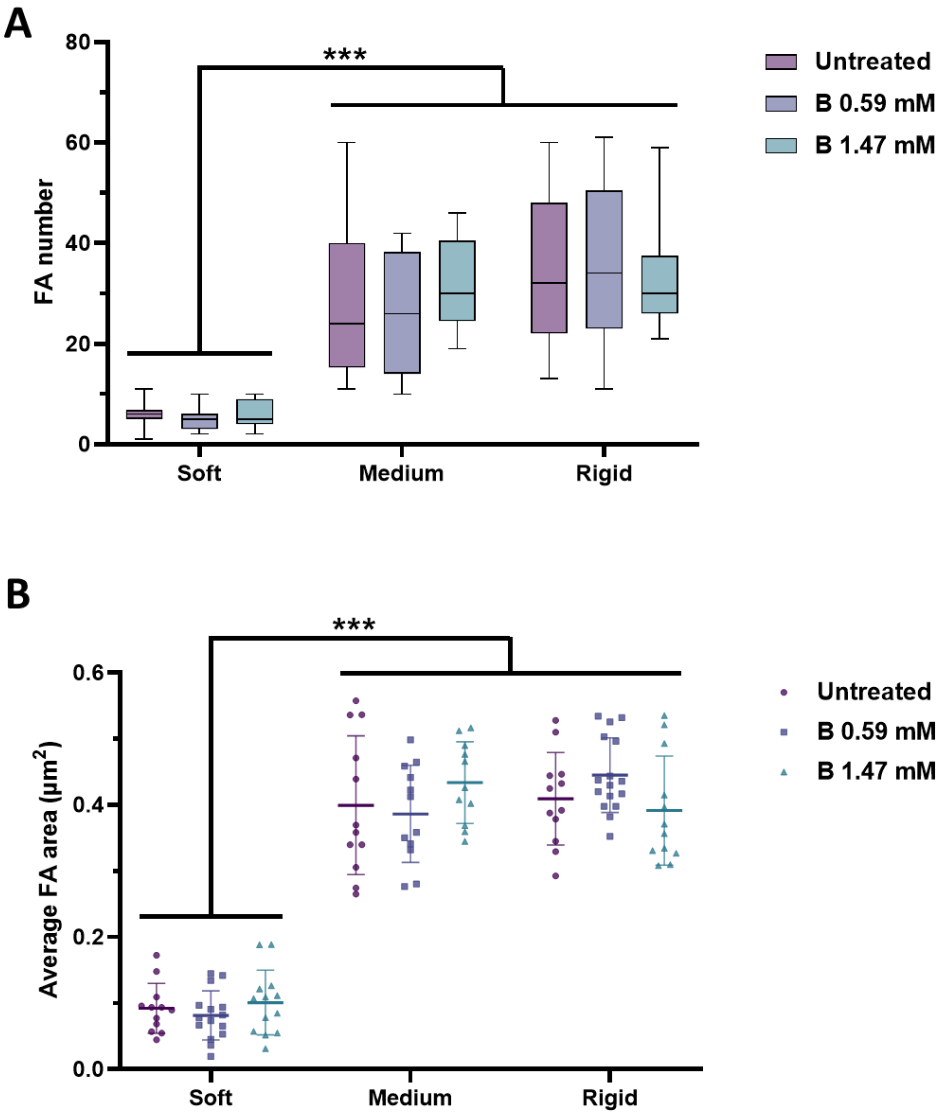


**Figure S8. The formation of FA in myoblasts on laminin-111 is not enhanced by NaBC1 stimulation.** Quantification of the number (A) and average area (B) of focal adhesions in C2C12 myoblasts seeded on PAAm hydrogels with different stiffness functionalised with laminin-111 and stimulated with soluble boron ions (0.59 and 1.47 mM). *n* = 10 cells from 3 different biological replicates. Data are represented as Mean ± Standard Deviation, and differences are considered significant for $p \leq 0.05$ using two-way ANOVAs (Tukey's multiple comparisons tests) for multiple comparisons. ***$p \leq 0.001$

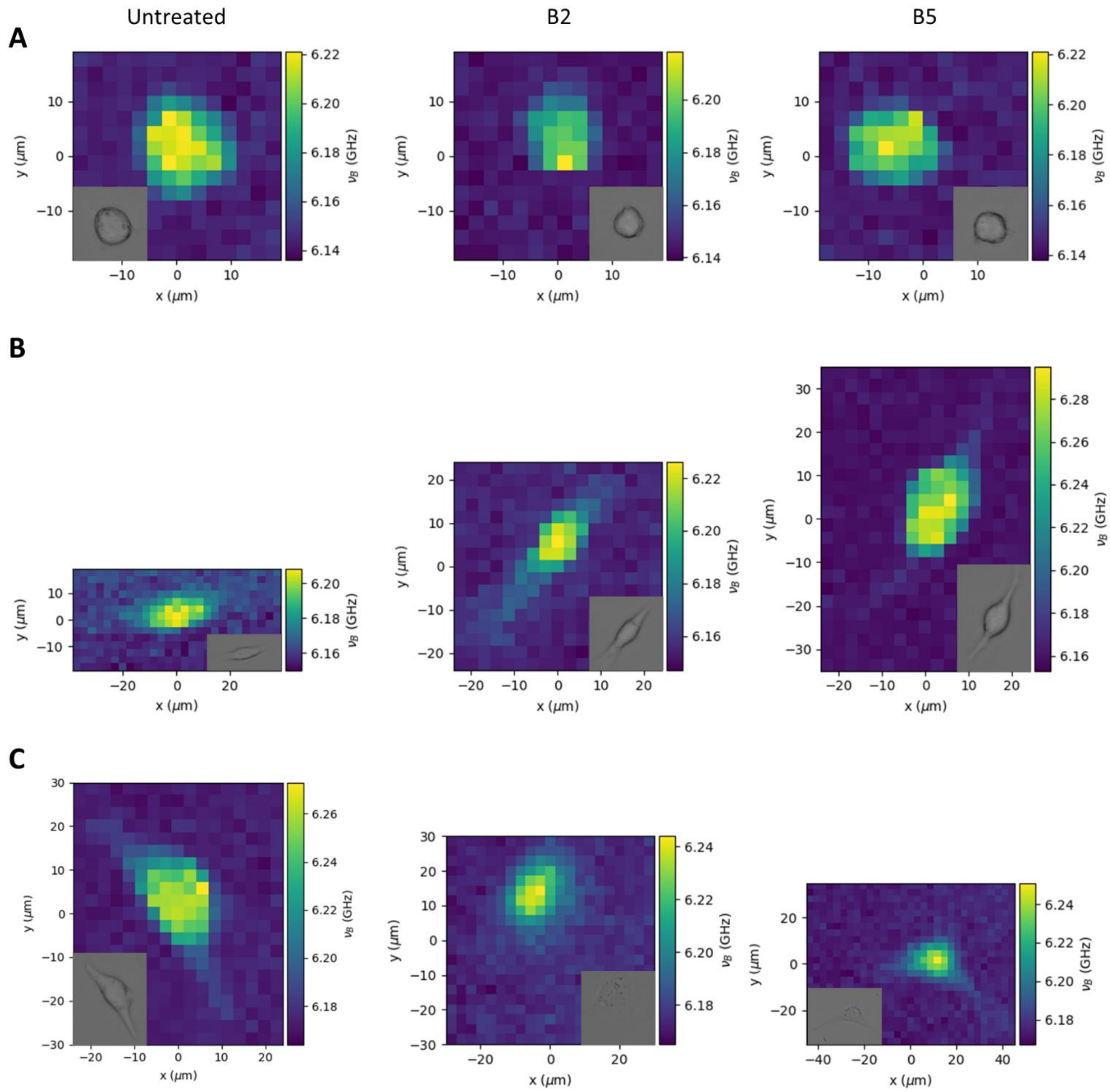


**Figure S9. Cell stiffness is not altered by NaBC1 stimulation on laminin-111.** Representative Brillouin maps of C2C12 myoblasts seeded on PAAm hydrogels with different stiffness functionalised with laminin-111 and stimulated with soluble boron (0.59 and 1.47 mM).

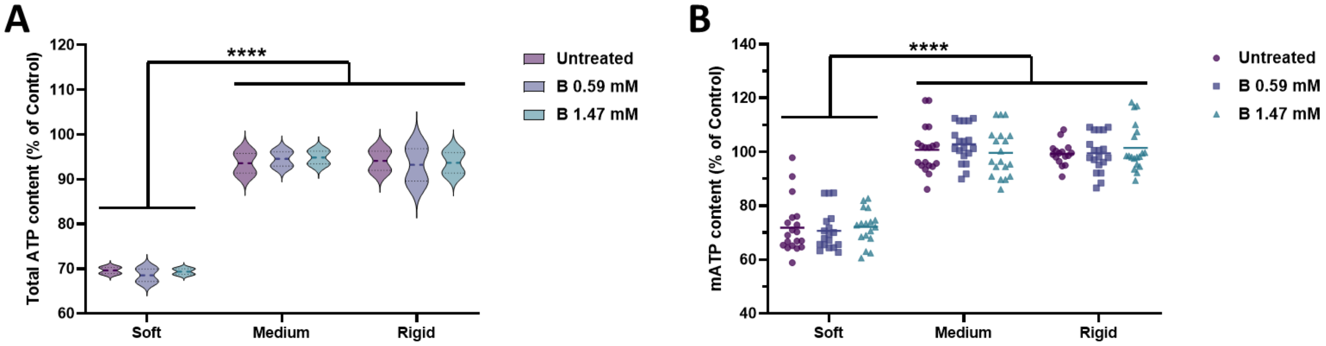


**Figure S10. Total and mitochondrial ATP content on laminin-111 is not influenced by NaBC1 or substrate stiffness.** A: Quantification of total content of C2C12 myoblasts seeded on PAAm hydrogels with different stiffness functionalised with laminin-111 and stimulated with soluble boron (0.59 and 1.47 mM). *n*: 3 biological replicates with 3 technical replicates. B: Quantification of mitochondrial content of C2C12 myoblasts seeded on PAAm hydrogels with different stiffness functionalised with laminin-111 and stimulated with soluble boron (0.59 and 1.47 mM). *n*: at least 10 cells from 3 biological replicates. Data are represented as Mean ± Standard Deviation, and differences are considered significant for $p \leq 0.05$ using two-way ANOVAs (Tukey's multiple comparisons tests) for multiple comparisons. ****$p \leq 0.0001$

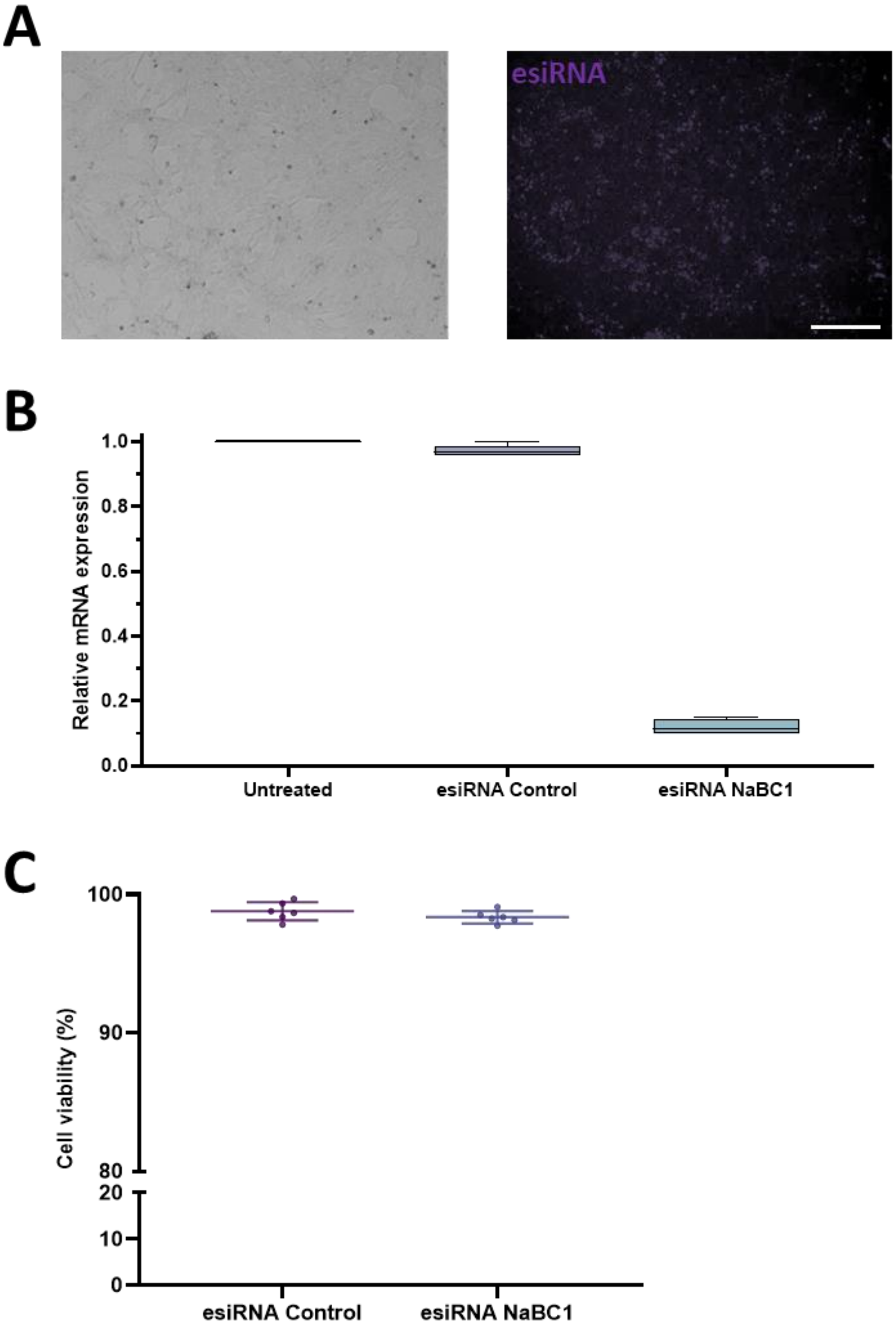


**Figure S11. NaBC1 silencing of C2C12 myoblasts does not affect cell viability.** A: Representative images of silenced NaBC1 C2C12 myoblasts. Magenta: Control esiRNA. Scale bar: 100 µm. B: Quantification of mRNA expression of NaBC1 in wild-type and silenced NaBC1 C2C12 myoblasts. *n*: 3 biological replicates with 3 technical replicates. C: Quantification of cell viability of silenced NaBC1 C2C12 myoblasts. *n*: 3 biological replicates with 3 technical replicates. Data are represented as Mean ± Standard Deviation, and differences are considered significant for $p \leq 0.05$ using one-way ANOVAs (Tukey's multiple comparisons tests) or *t*-tests for multiple or pairwise comparisons, respectively. ****$p \leq 0.0001$

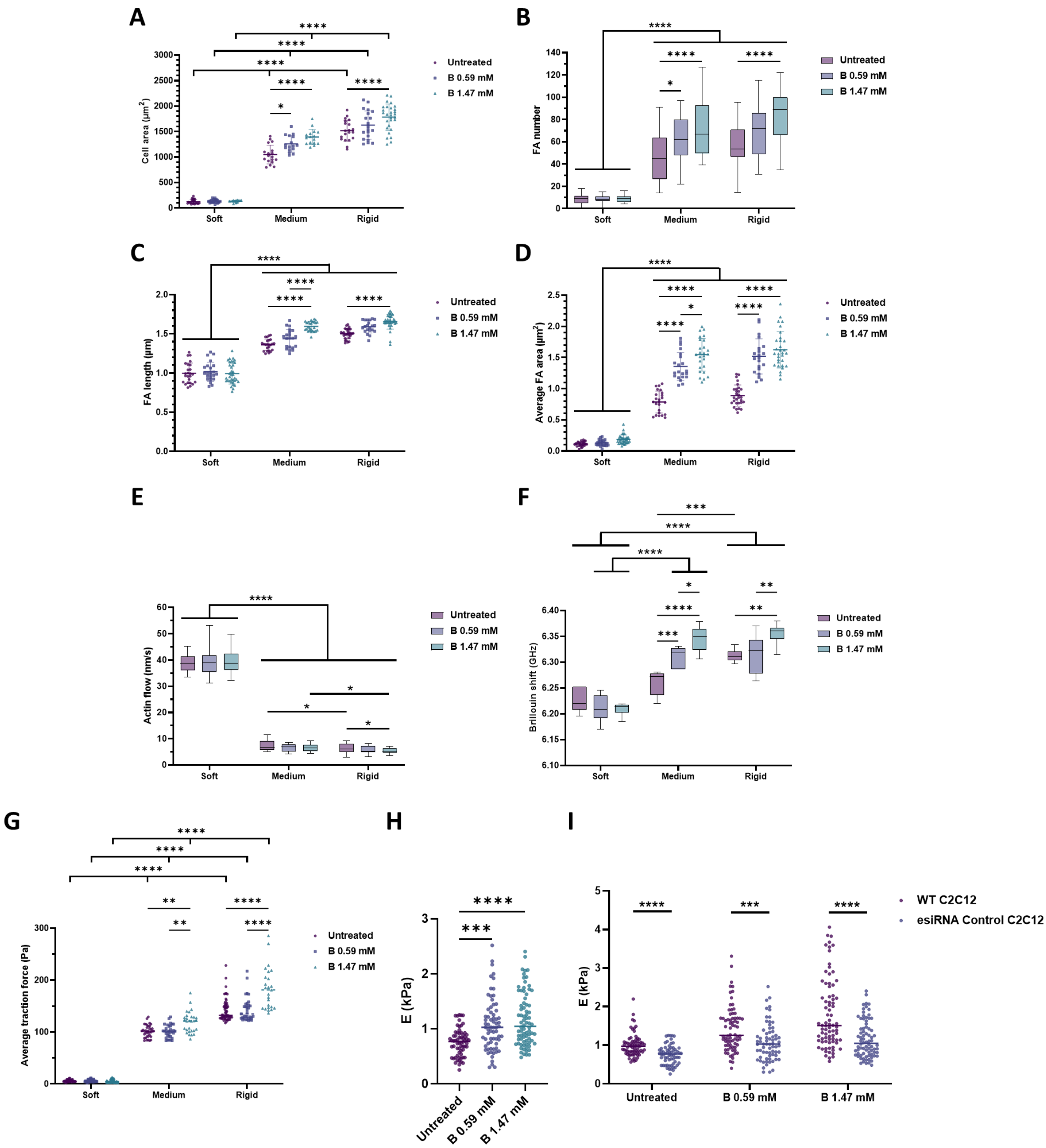


**Figure S12. Transfection of C2C12 myoblasts does not affect cell behavior.** The results reported in panels A-D derive from experiments in which C2C12 myoblasts were transfected with esiRNA Control and seeded on PAAm hydrogels of different stiffnesses (soft, medium, and rigid) that were functionalized with fibronectin (FN) and stimulated with soluble boron ions (B) at two different concentrations (0.59 and 1.47 mM). A: Quantification of cell area of Control-silenced C2C12 myoblasts that were treated and cultured as described. *n* = 10 cells from 3 different biological replicates. B: Quantification of the number of focal adhesions in Control-silenced C2C12 myoblasts seeded on PAAm hydrogels with different stiffness functionalised with fibronectin and stimulated with soluble boron (0.59 and 1.47 mM). *n* = 10 cells from 3 different biological replicates. C: Quantification of focal adhesion (FA) length in Control-silenced C2C12 myoblasts that were treated and cultured as described. *n* = 10 cells from 3 different biological replicates. D: Quantification of focal adhesion (FA) average area in Control-silenced C2C12 myoblasts that were treated and cultured as described. *n* = 10 cells from 3 different biological replicates. E: Quantification of actin retrograde flow in Control-silenced C2C12 myoblasts that were treated and cultured as described. *n* = 5 cells with at least 5 different flow areas per cell. F: Quantification of Brillouin shift in Control-silenced C2C12 myoblasts that were treated and cultured as described and imaged by Brillouin microscopy. *n* = 10 cells from 3 different biological replicates. G: Quantification of traction forces exerted by Control-silenced C2C12 myoblasts that were treated and cultured as described. *n* = 30 cells from 10 different locations within each hydrogel from 3 different biological replicates. H: Quantification of cell stiffness by nanoindentation of Control-silenced C2C12 myoblasts seeded on glass coverslips functionalized with FN and stimulated with soluble B (0.59 and 1.47 mM). *n* = 10 cells with 9 indentations on each single cell from 3 different biological replicates. I: Comparison of cell stiffness by nanoindentation of wild type and Control-silenced C2C12 myoblasts seeded on glass coverslips functionalized with FN and stimulated with soluble B (0.59 and 1.47 mM). *n* = 10 cells with 9 indentations on each single cell from 3 different biological replicates. Data are represented as Mean ± Standard Deviation, and differences are considered significant for $p \leq 0.05$ using one-way ANOVAs or two-way ANOVAs (Tukey's multiple comparisons tests) for multiple comparisons. *$p \leq 0.05$, **$p \leq 0.01$, ***$p \leq 0.001$, ****$p \leq 0.0001$

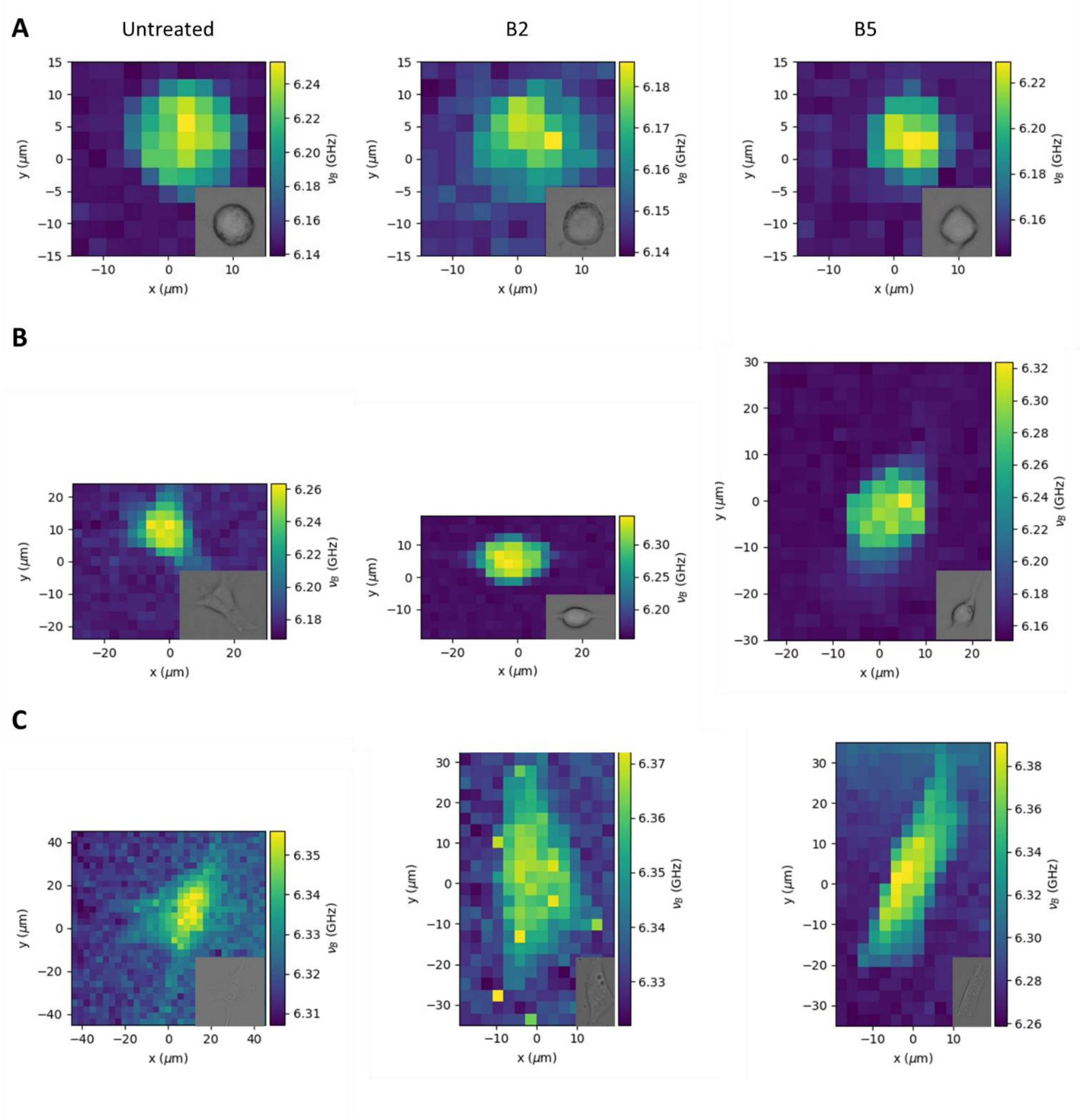


**Figure S13. Control-silencing does not alter cell stiffness on fibronectin-coated substrates.** Representative Brillouin maps of Control-silenced C2C12 myoblasts seeded on PAAm hydrogels with different stiffness functionalised with fibronectin and stimulated with soluble boron (0.59 and 1.47 mM).

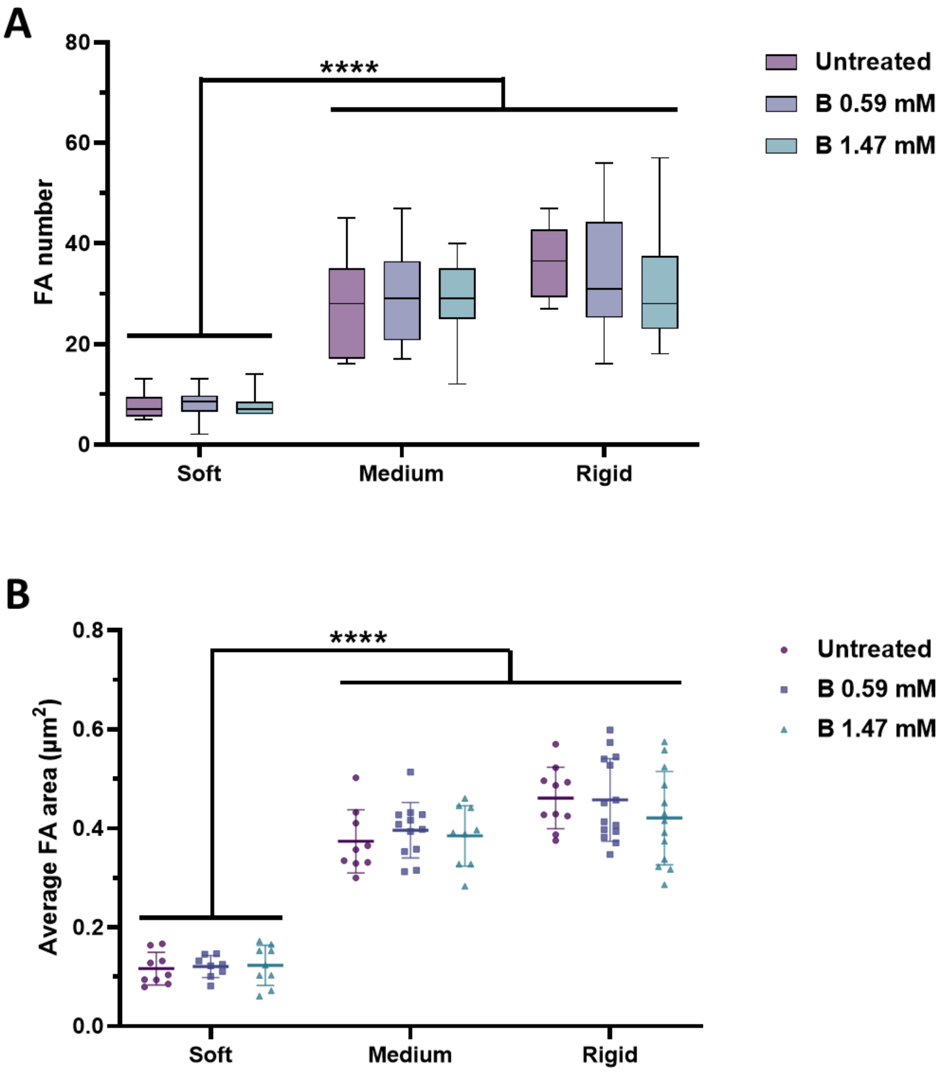


**Figure S14. NaBC1 regulates the stiffness-mediated triggering of formation of FA on fibronectin in NaBC1-silenced myoblasts.** Quantification of the number (A) and average area (B) of focal adhesions in C2C12 myoblasts seeded on PAAm hydrogels with different stiffness functionalised with fibronectin and stimulated with soluble boron (0.59 and 1.47 mM). *n* = 10 cells from 3 different biological replicates. Data are represented as Mean ± Standard Deviation, and differences are considered significant for $p \leq 0.05$ using two-way ANOVAs (Tukey's multiple comparisons tests) for multiple comparisons. ****$p \leq 0.0001$

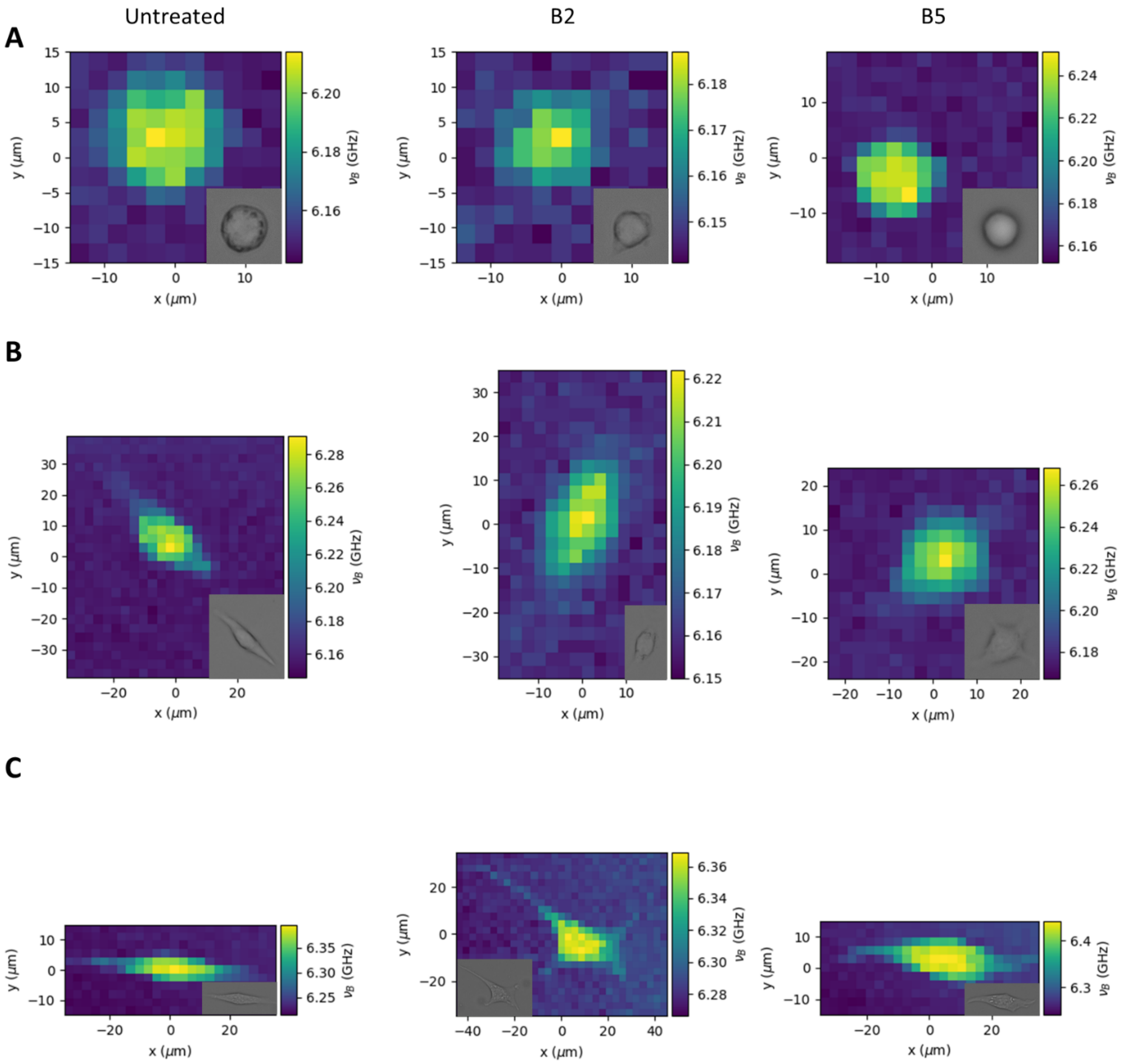


**Figure S15. NaBC1-silencing does not alter cell stiffness on fibronectin-coated substrates.** Representative Brillouin maps of NaBC1-silenced C2C12 myoblasts seeded on PAAm hydrogels with different stiffness functionalised with fibronectin and stimulated with soluble boron (0.59 and 1.47 mM).

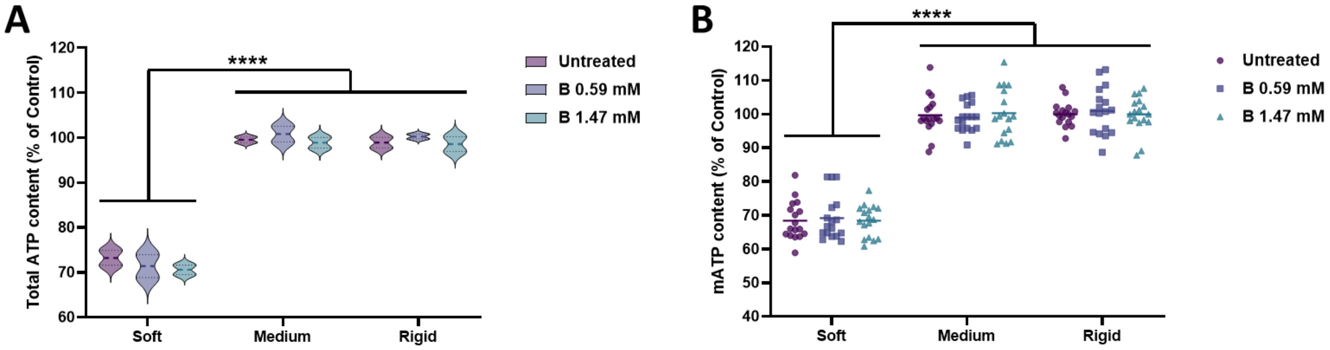


**Figure S16. NaBC1 stimulation is essential for stiffness-mediated triggering of total and mitochondrial ATP content on fibronectin substrates.** A: Quantification of total content of NaBC1-silenced C2C12 myoblasts seeded on PAAm hydrogels with different stiffness functionalised with fibronectin and stimulated with soluble boron (0.59 and 1.47 mM). *n*: 3 biological replicates with 3 technical replicates. B: Quantification of mitochondrial content of NaBC1-silenced C2C12 myoblasts seeded on PAAm hydrogels with different stiffness functionalised with fibronectin and stimulated with soluble boron (0.59 and 1.47 mM). *n*: at least 10 cells from 3 biological replicates. Data are represented as Mean ± Standard Deviation, and differences are considered significant for $p \leq 0.05$ using two-way ANOVAs (Tukey's multiple comparisons tests) for multiple comparisons. ****$p \leq 0.0001$

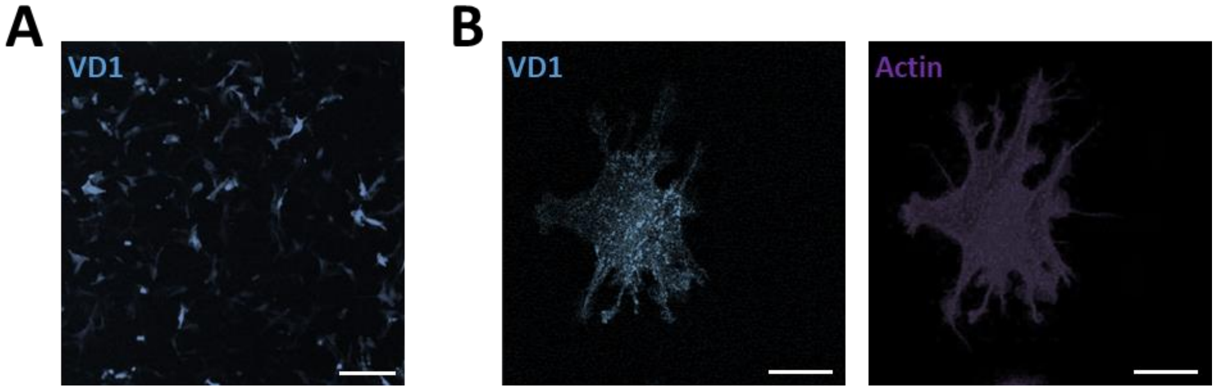


**Figure S17. Transfection of C2C12 myoblasts with the VD1 plasmid.** A: Representative image of C2C12 myoblasts transfected with the VD1 plasmid. Cyan: VD1 plasmid. Scale bar: 100 µm. B: Representative images of C2C12 myoblasts transfected with the VD1 and LifeAct plasmids. Cyan: VD1 plasmid; Magenta: LifeAct plasmid. Scale bar: 20 µm.

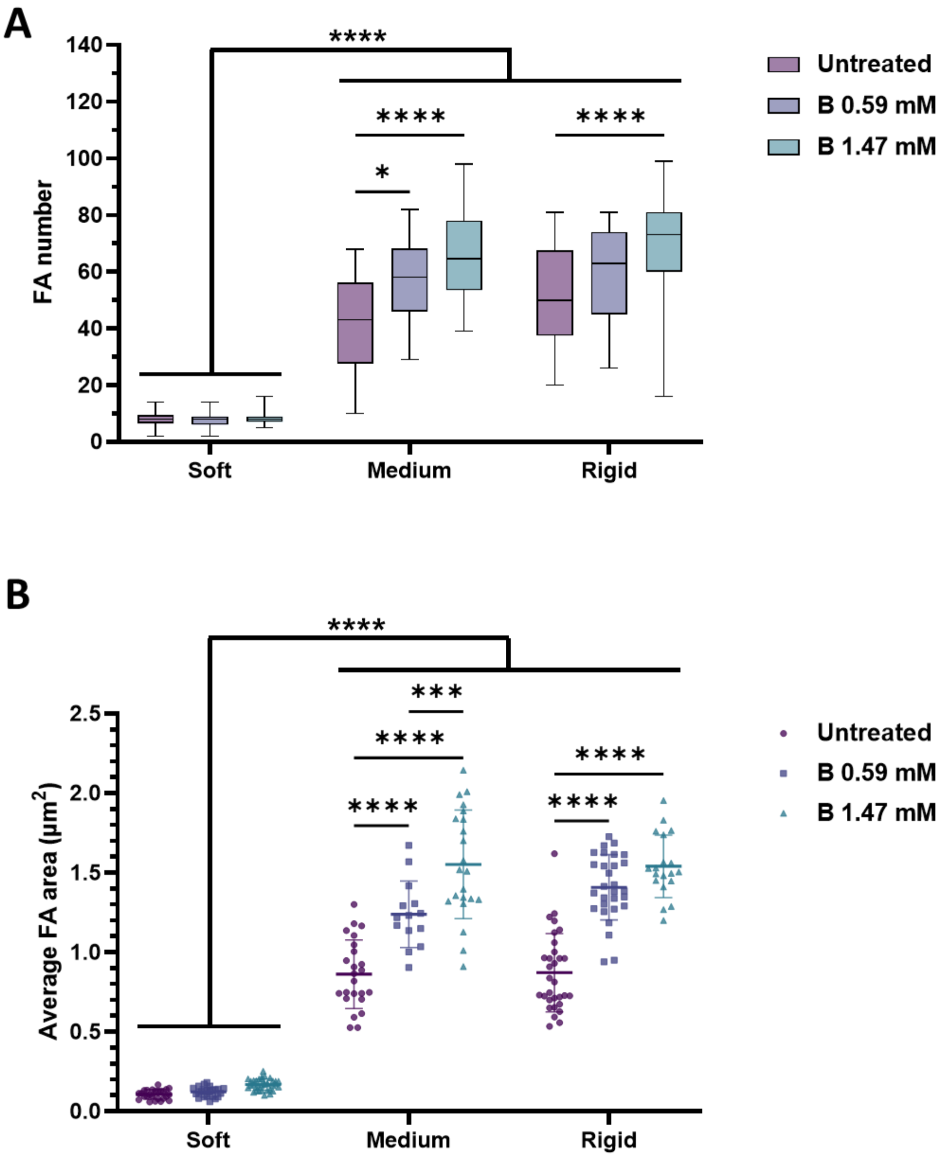


**Figure S18. NaBC1 and stiffness-mediated triggering of formation of FA in myoblasts on fibronectin substrates is independent of the talin-vinculin binding.** Quantification of the number (A) and average area (B) of focal adhesions in C2C12 myoblasts transfected with the VD1 plasmid seeded on PAAm hydrogels with different stiffness functionalised with fibronectin and stimulated with soluble boron (0.59 and 1.47 mM). *n* = 10 cells from 3 different biological replicates. Data are represented as Mean ± Standard Deviation, and differences are considered significant for $p \leq 0.05$ using two-way ANOVAs (Tukey's multiple comparisons tests) for multiple comparisons. *$p \leq 0.05$, ****$p \leq 0.0001$

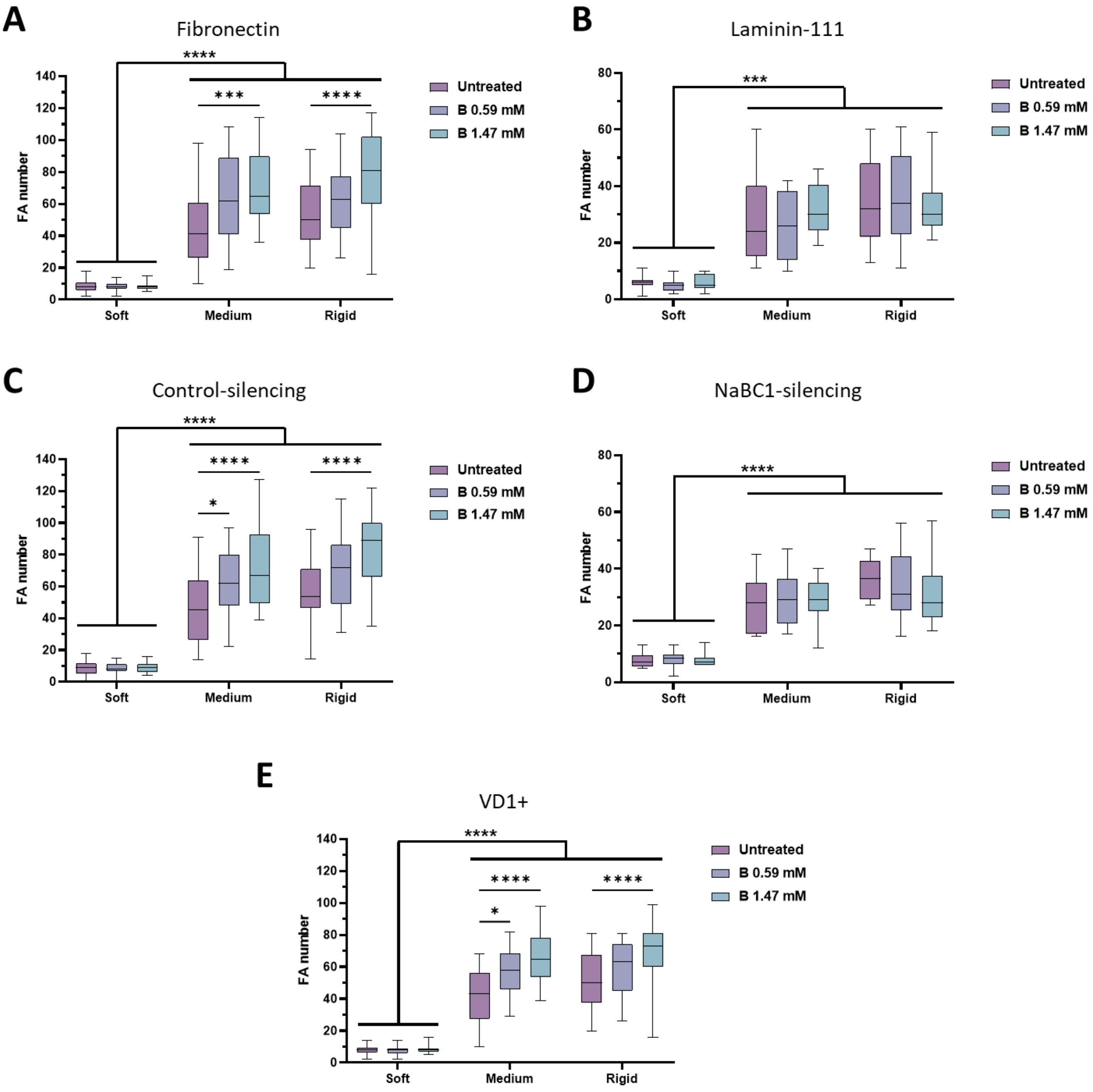


**Figure S19. Comparison of the number of FA in C2C12 myoblasts.** Quantification of the number of focal adhesions in wild type C2C12 myoblasts (A-B), Control-silenced C2C12 myoblasts (C), NaBC1-silenced C2C12 myoblasts (D) or C2C12 myoblasts transfected with the VD1 plasmid (E) seeded on PAAm hydrogels with different stiffness functionalised with fibronectin (A, C, D and E) or laminin-111 (B) and stimulated with soluble boron (0.59 and 1.47 mM). *n* = 10 cells from 3 different biological replicates. Data are represented as Mean ± Standard Deviation, and differences are considered significant for $p \leq 0.05$ using two-way ANOVAs (Tukey's multiple comparisons tests) for multiple comparisons. *$p \leq 0.05$, ***$p \leq 0.001$, ****$p \leq 0.0001$

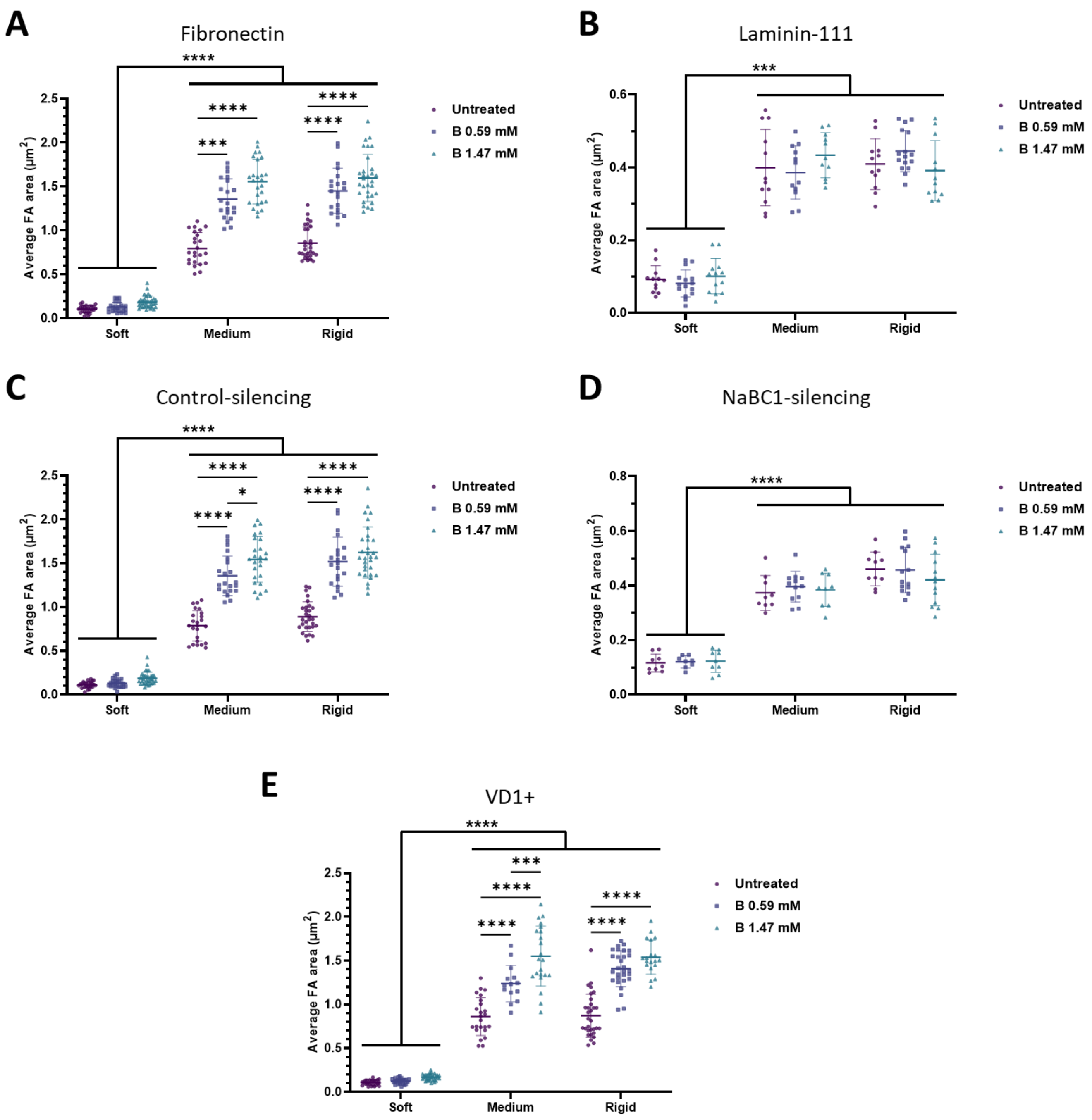


**Figure S20. Comparison of the average FA area in C2C12 myoblasts.** Quantification of the average area of focal adhesions in wild type C2C12 myoblasts (A-B), Control-silenced C2C12 myoblasts (C), NaBC1-silenced C2C12 myoblasts (D) or C2C12 myoblasts transfected with the VD1 plasmid (E) seeded on PAAm hydrogels with different stiffness functionalised with fibronectin (A, C, D and E) or laminin-111 (B) and stimulated with soluble boron (0.59 and 1.47 mM). *n* = 10 cells from 3 different biological replicates. Data are represented as Mean ± Standard Deviation, and differences are considered significant for $p \leq 0.05$ using two-way ANOVAs (Tukey's multiple comparisons tests) for multiple comparisons. *$p \leq 0.05$, ***$p \leq 0.001$, ****$p \leq 0.0001$

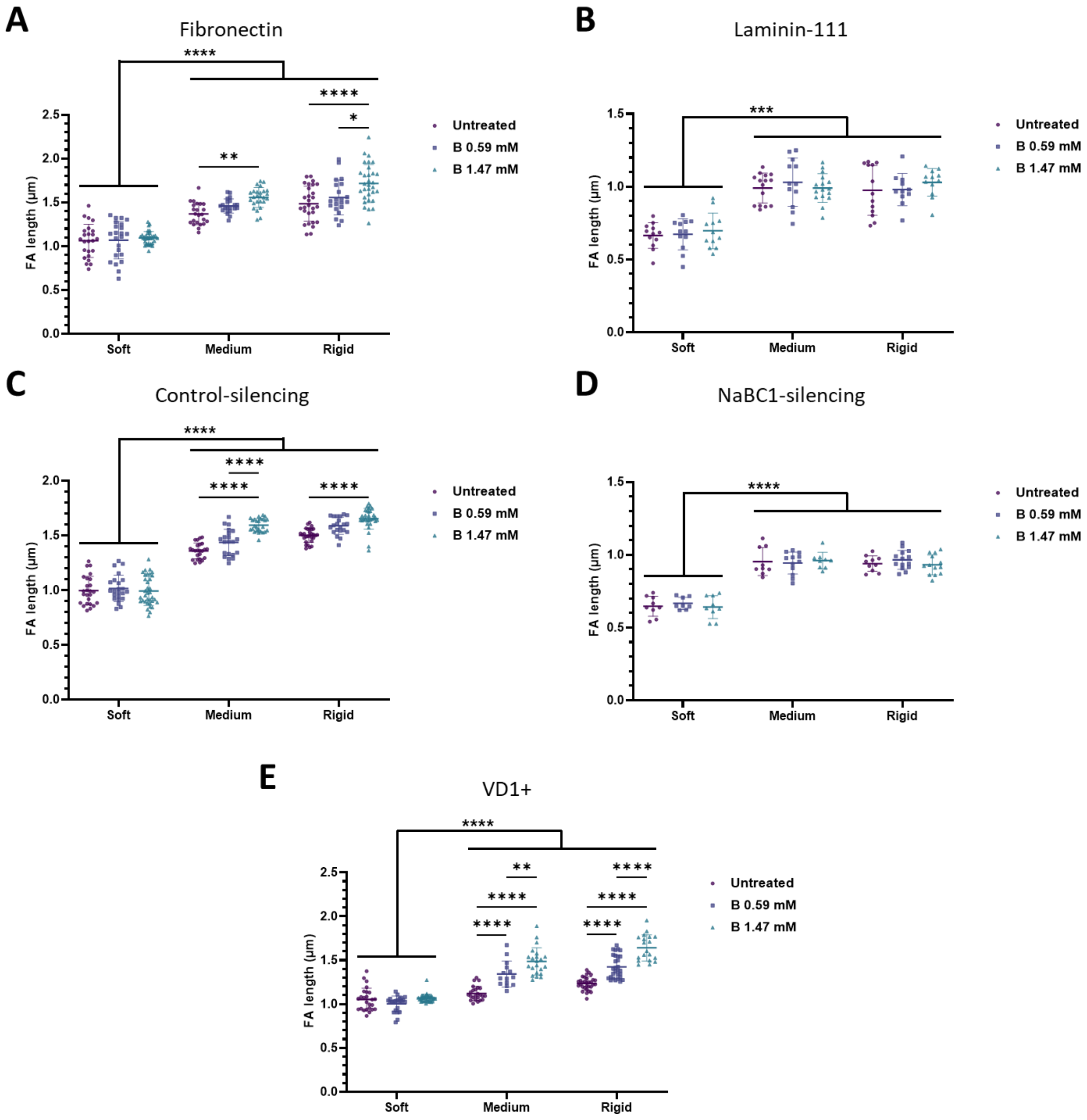


**Figure S21. Comparison of FA length in C2C12 myoblasts.** Quantification of the length of focal adhesions in wild type C2C12 myoblasts (A-B), Control-silenced C2C12 myoblasts (C), NaBC1-silenced C2C12 myoblasts (D) or C2C12 myoblasts transfected with the VD1 plasmid (E) seeded on PAAm hydrogels with different stiffness functionalised with fibronectin (A, C, D and E) or laminin-111 (B) and stimulated with soluble boron (0.59 and 1.47 mM). *n* = 10 cells from 3 different biological replicates. Data are represented as Mean ± Standard Deviation, and differences are considered significant for $p \leq 0.05$ using two-way ANOVAs (Tukey's multiple comparisons tests) for multiple comparisons. *$p \leq 0.05$, **$p \leq 0.01$, ***$p \leq 0.001$, ****$p \leq 0.0001$
